# Structural basis for synthase activation and cellulose modification in the *E. coli* Type II Bcs secretion system

**DOI:** 10.1101/2024.06.05.597511

**Authors:** Itxaso Anso, Samira Zouhir, Thibault G. Sana, Petya Violinova Krasteva

## Abstract

Bacterial cellulosic polymers constitute a prevalent class of biofilm matrix exopolysaccharides that rely on conserved cyclic diguanylate (c-di-GMP)-dependent cellulose synthases. Polymer structure and modifications, however, depend on the ensemble of synthase modules and accessory subunits, thus defining several types of bacterial cellulose secretion (Bcs) systems. In *E. coli*, a BcsRQABEFG macrocomplex, encompassing the inner membrane and cytosolic subunits, and an outer membrane porin, BcsC, secure the biogenesis of phosphoethanolamine (pEtN)-modified cellulose. Resolution-limited studies have proposed different macrocomplex stoichiometries and its assembly and regulation have remained elusive. Using cryo-EM, we visualize the molecular mechanisms of BcsA-dependent recruitment and stabilization of a trimeric BcsG pEtN-transferase for polymer modification and a dimeric BcsF-dependent recruitment of an otherwise cytosolic BcsE_2_R_2_Q_2_ regulatory complex. We further demonstrate that BcsE, a secondary c-di-GMP sensor, remains dinucleotide-bound and retains the essential-for-secretion BcsRQ partners onto the synthase even in the absence of direct c-di-GMP-synthase complexation, likely lowering the threshold for c-di-GMP-dependent synthase activation. Such ‘activation-by-proxy’ mechanism could allow Bcs secretion system activation even in the absence of dramatic intracellular c-di-GMP increase and is reminiscent of other widespread synthase-dependent polysaccharide secretion systems where c-di-GMP sensing and/or synthase stabilization are carried out by key co-polymerase subunits.

## Introduction

Bacteria have evolved sophisticated nanomachines for the biogenesis of extracellular biofilm matrix components, which allow them to achieve cooperative multicellularity, increased fitness and homeostasis ^1–4^. Across the bacterial domain of life, and especially in Gram-negative pathogens and eukaryotic host-associated microbes such as *E. coli* or *S. enterica* serovar Typhimurium, biofilm formation is typically controlled by the RNA-based second messenger c-di-GMP, which is able to elicit multiple pathway-specific physiological responses via spatially restrained intracellular signaling mechanisms^5^. Bacterial synthase-dependent exopolysaccharide secretion systems are prevalent c-di-GMP sensor-effectors, which incorporate modules for dinucleotide sensing, glycan polymerization, transmembrane export, synthase regulation and polymer modification, and thus determine the physicochemical properties of the mature biofilms and their interactions with environment and/or eukaryotic hosts^4,6^.

Bacterial cellulosic polymers are a widespread class of biofilm matrix exopolysaccharides that find a variety of biotechnological applications and are synthesized by dedicated cellulose synthase enzymes^1,6^. The latter’s core fold of a glycosyl transferase (GT) and an membrane transport (TMD) domains is structurally conserved across kingdoms^6,7^, however the bacterial BcsA orthologs typically incorporate an additional PilZ domain for c-di-GMP-dependent synthase regulation^8,9^. Nevertheless, cellulose secretion and the actual polymer structure and modifications are determined not only by the ensemble of synthase modules, but also by a multitude of accessory subunits, which can assemble in several distinct types of Bcs secretion systems^4,6,10^. In particular, type I Bcs systems are characterized by the expression of BcsD proposed to engage in a variety of intracellular scaffolds for both crystalline and modified cellulose secretion, type II systems feature the BcsE and BcsG components discussed below, and type III systems lack all BcsD, BcsE and BcsG subunits and often feature BcsK (periplasmic scaffolding only) rather than BcsC (scaffolding and outer membrane export) homologs in the periplasm^4,10^.

In *E. coli* and other enterobacteria, the biofilm matrix is composed primarily of proteinaceous fimbriae, such as non-motile flagella and amyloid curli, and of cellulosic polymers produced by an *E. coli*-like or type II Bcs secretion system, which incorporates additional c-di-GMP-sensing and polymer modification modules^3,6^ (Fig. 1a). In particular, *E. coli* cellulose biogenesis requires the concerted expression of two adjacent *bcs* operons (*bcsRQABZC* and *bcsEFG*), whose protein products assemble into a multicomponent BcsRQABEFG synthase macrocomplex embedded in the inner membrane, a periplasmic cellulase (BcsZ) and an outer membrane porin with periplasmic scaffolding repeats (BcsC)^11^ (Fig. 1a). Whereas *in vitro* cellulose synthesis can be carried out with only the BcsA synthase and a C-terminal tail anchor (TA) from the co-polymerase BcsB, micromolar concentrations of activating c-di-GMP, bivalent ions (e.g. Mg^++^) and uridine diphosphate glucose (UDP-glucose) as energetically preloaded substrate^12^, the rest of the Bcs macrocomplex components are either essential or act as enhancers for cellulose biosynthesis *in vivo*^11^. Of these, the BcsG subunit has been shown to interact with a *E. coli* type-specific N-terminal domain of the BcsA synthase^11^ and to introduce phosphoethanolamine (pEtN) moieties onto the nascent polymer via a pEtN-transferase domain in the periplasm^13^; the transmembrane peptide BcsF has been shown to recruit the secondary c-di-GMP-sensing protein BcsE^14^; and the latter - together with an essential-for-secretion BcsRQ ATPase complex - has been shown to form a cytosolic vestibule around the synthase’s cytosolic modules^11,15^ (Fig. 1a). Whereas fragmentary insights into Bcs macrocomplex formation and components’ structures have been obtained from several crystallographic and electron microscopy studies^11,14–16^, the overall stoichiometry, assembly and regulatory mechanisms have remained enigmatic.

**Fig. 1.**
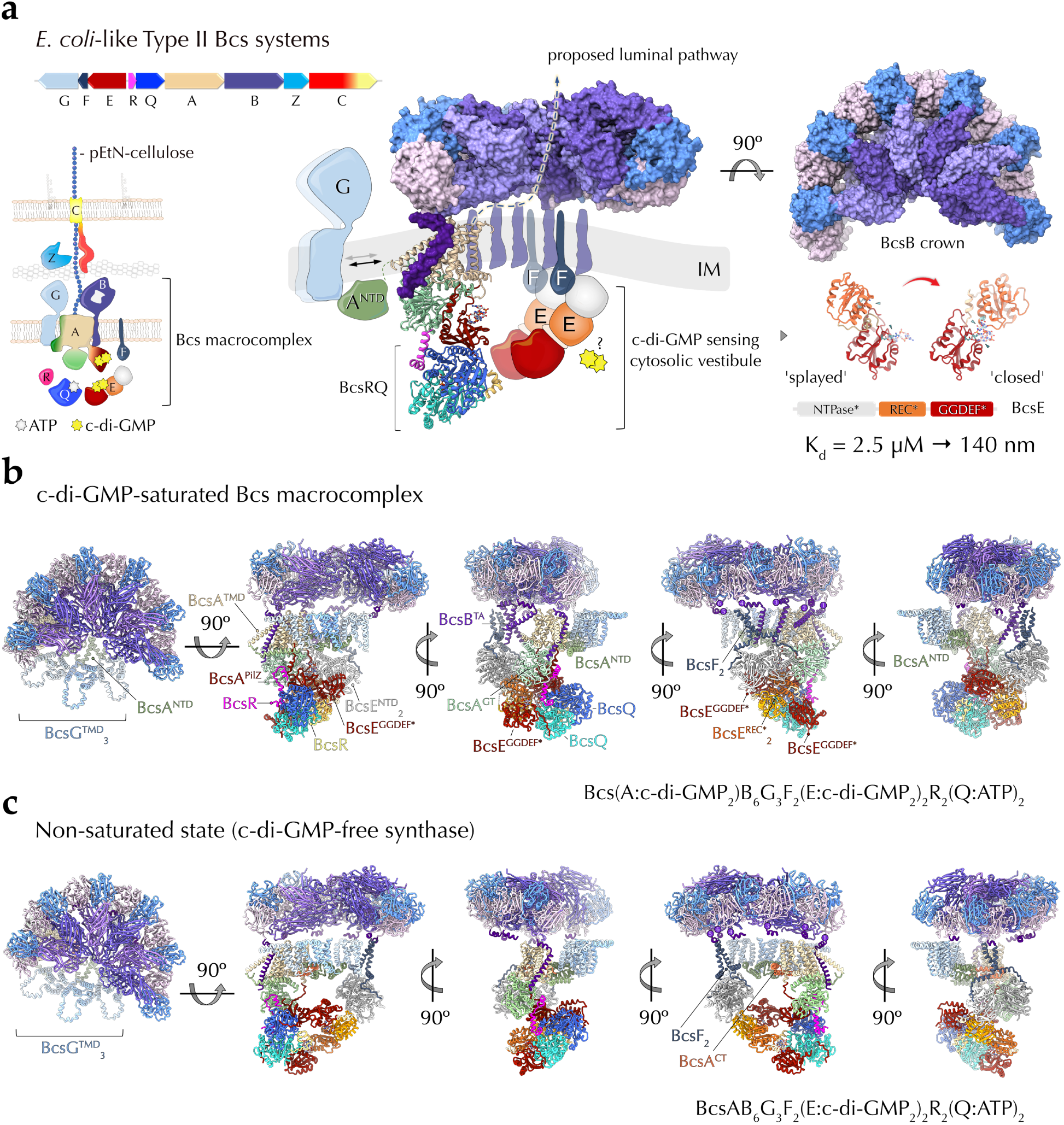
State-of-the-art and here-in presented structures of the Bcs macrocomplex from the E. coli Type II cellulose secretion system. **a** Left, *E. coli bcs* operon organization and thumbnail representation of the secretion system topology in the *E. coli* envelope. Right, current structural insights into complex assembly from X-ray crystallographic and electron microscopy structures^11,14–16^. Densities for BcsA^NTD^, BcsG, BcsE and BcsF have remained largely unresolved in the, whereas crystallographic snapshots have captured two different conformations of BcsE^14,15^. The formation of a composite c-di-GMP binding site by RxxD motifs from both the degenerate REC* and GGDED* domains increases the affinity for dimeric c-di-GMP from the low micromolar to nanomolar range^15^. c-di-GMP, cyclic diguanylate; pEtN, phosphoethanolamine; NTD, N-terminal domain; IM, inner membrane; REC*, phosphorylation-incompetent receiver domain; NTPase*, catalytically incompetent nucleoside triphosphatase domain; GGDEF*, degenerate diguanylate cyclase domain. **b** Cartoon representation of the here-in resolved cryo-EM structure of the assembled c-di-GMP-saturated Bcs macrocomplex in five different views. TMD, transmembrane domain; GT, glycosyl transferase domain; TA, tail-anchor; ATP, adenosine triphosphate. c Cartoon representation of the here-in resolved cryo-EM structure of the assembled Bcs macrocomplex, featuring a c-di-GMP-free BcsA. CT, C-terminal amphipathic helices.

Here we use cryogenic electron microscopy (cryo-EM) to settle conflicting reports on the macrocomplex stoichiometry^15,16^ and reveal the molecular mechanisms of regulatory subunit recruitment and function. We demonstrate that BcsA’s N-terminal domain adopts an amphipathic fold to recruit three copies of the pEtN transferase BcsG, stabilized opposite of the previously reported hexameric BcsB crown. We further demonstrate that the single-pass inner membrane polypeptide BcsF, which has evaded earlier structural characterization, folds into an X-shaped dimer whose extended cytosolic tails recruit two copies of BcsE via the latter’s N-terminal domains in a cytosolic β-sheet complementation mechanism. These interactions are also necessary to recruit and retain the regulatory BcsR_2_Q_2_ complex onto the synthase’s cytosolic modules, where the key for secretion N-terminal domain of BcsR plugs into a hydrophobic pocket on the BcsA^GT^ domain and together with BcsQ stabilizes the relative orientation of the adjacent PilZ module. This regulatory mechanism is likely conserved across not only *E. coli*-like type II Bcs secretion systems in γ-proteobacterial *Enterobacteriaceae*, but also in type I Bcs systems across the β-proteobacterial clade featuring BcsPD cytoskeletal scaffolds for cellulose secretion regulation^17^. Finally, by Bcs macrocomplex purification at dinucleotide concentrations close to the dissociation constant for BcsA-c-di-GMP complexation, we visualize both the dinucleotide-free and dinucleotide-bound states for the synthase (Fig. 1b-c and Supplementary Fig. 1). Importantly, the higher-affinity BcsE sensor remains c-di-GMP-bound in both structures and retains a compact conformation^15^ to retain the regulatory BcsR_2_Q_2_ complex onto the synthase, likely preserving the latter’s catalytically competent state. Together, these data suggest that by the recruitment of the multicomponent BcsE_2_R_2_Q_2_ vestibule around the BcsA^PilZ^ domain, the *E. coli*-like Bcs macrocomplex has likely evolved a cooperative ‘activation-by-proxy’ mechanism to lower the threshold for c-di-GMP-dependent activation and to secure cellulose secretion initiation even in the early biofilm-forming stages and/or layers that are characterized by low intracellular c-di-GMP and the presence of counteracting phosphodiesterases^3^.

## Results

### Stoichiometry of the assembled Bcs macrocomplex of E. coli

Bacterial BcsA orthologs are processive glycosyl transferase 2 family synthases with a single cytosolic GT domain that uses UDP-glucose as substrate in an inverting, divalent metal ion-dependent mechanism of glycan polymerization, best studied *in vitro* in the *Rhodobacter sphaeroides* BcsAB heterodimeric complex^9,18,19^. Polymerization is coupled with inner membrane polysaccharide extrusion through a narrow pore in the BcsA^TMD^-BcsB^TA^ inner membrane complex translocating a non-modified, non-hydrated homopolymer. In the resting state the BcsA^GT^ active site is capped by a so-called ‘gating loop’, stabilized by interactions with the N-proximal BcsA^PilZ^ domain linker that senses c-di-GMP^18^. In the presence of micromolar concentrations of dinucleotide the PilZ domain undergoes an approximately 18° rotation and 4.4 Å displacement around a C-proximal α−helical ‘hinge’ and the linker-gating loop stabilizing interactions are released to yield a catalytically competent state^9^. Processive cycles of active site opening, substrate entry, gating loop closure, polymerization and translocation are then determined by the presence of product vs. substrate in the active site and minute movements of a so-called ‘finger helix’ in the bottom of the enzyme’s active site^6,19^.

Whereas BcsA itself is highly conserved, the secretion-competent synthase macrocomplexes are strikingly diverse across the bacterial clade^4,6^. In *E. coli*, in particular, the catalytic BcsAB tandem has been shown the associate with the ensemble of inner membrane and cytosolic subunits in an approximately megadalton-sized secretory assembly. In it, the synthase associates in a non-canonical BcsA:BcsB stoichiometry with up to six BcsB copies whose donut-shaped periplasmic modules assemble into a superhelical crown with stacking carbohydrate-binding modules likely guiding the extruded polysaccharide into the periplasm and towards the BcsC periplasmic scaffold^11,15^. Additionally, the synthase has been found to associate stably with an essential-for-secretion BcsRQ tandem, with the secondary c-di-GMP sensing protein BcsE, the inner membrane polypeptide BcsF and the pEtN-transferase BcsG^11,15,16^ (Fig. 1a). The stoichiometry and recruitment mechanisms of all of these latter components have remained under debate, mostly due to the limited resolutions of previously reported structural models in the literature^11,15,16^.

Here we present cryo-EM structures of the *E. coli* Bcs macrocomplex positioning all seven BcsRQABEFG partners and multiple c-di-GMP-binding sites (Fig. 1b-c). We show that the Bcs macrocomplex contains a single synthase associated with a hexameric BcsB crown on one side and a trimeric BcsG pEtN-transferase complex recruited by the *E. coli*-specific BcsA^NTD^ on the other. We further show that the BcsA cytosolic domains interact with a heterotetrameric BcsR_2_Q_2_ complex in the cytosol. The latter further is retained by a dimeric BcsE tandem, whose degenerate receiver (REC*) and diguanylate cyclase (GGDEF*) domains bind an intercalated dimeric c-di-GMP per BcsE protomer. The previously uncharacterized N-terminal domains (NTD) of BcsE, on the other hand, form a membrane proximal head-to-head dimer and adopt a P-loop nucleotide triphosphatase (NTPase)-like fold whose central β-sheet is extended by the C-terminal tails of an inner membrane-embedded BcsF dimer in proximity to the tail anchors of the synthase-distal BcsB copies from the crown. Together, these results demonstrate a definitive BcsR_2_Q_2_AB_6_E_2_F_2_G_3_ stoichiometry for the assembled *E. coli* Bcs macrocomplex, which binds up to six c-di-GMP molecules for maximal synthase activation and nascent polysaccharide modification (Fig. 1b-c). Finally, we visualize the complex’s intrinsic conformational plasticity, in which the regulatory vestibule complex, and BcsE in particular, can resort to alternative protein interaction interfaces to maintain the activating BcsRQ partners onto the synthase cytosolic modules (Fig. 1b-c).

### The inner membrane BcsAB_6_G_3_F_2_ complex

We demonstrated previously that contrary to the canonical 1:1 BcsAB assemblies observed in purified samples from *Rhodobacter sphaeroides* (Type III Bcs secretion system)^18^ and *G. hansenii* (Type I Bcs secretion system)^20^, in the assembled *E. coli* Bcs macrocomplex BcsA associates with a BcsB hexamer whose periplasmic modules of alternating carbohydrate-binding and flavodoxin-like domains (CBD_1_-FD_1_-CBD_2_-FD_2_) polymerize via a β-sheet complementation mechanism between the FD_1_^n^:FD_2_^n+1^ domains of adjacent BcsB protomers^15^. The structure of the hexameric periplasmic crown is refined here to 2.35 Å resolution and reveals to near-atomic detail the molecular mechanism of BcsB polymerization, with more than 3300 Å^2^ interface surface and a free energy gain of −22.5 kcal/mol between each pair of adjacent protomers (Supplementary Fig. 2).

Whereas earlier studies have visualized the overall fold of the transmembrane regions of the BcsA synthase and the C-terminal tail-anchor of its associated BcsB protomer^15,16^, here we position the transmembrane anchors for most of the BcsB subunits and present the first structures of the regulatory BcsG and BcsF partners at side-chain resolution. The first and second copies BcsB copies engage in contacts with the BcsA synthase, whereas the synthase-distal BcsB copies position close to an X-shaped transmembrane BcsF dimer. In particular, the BcsF tandem positions between the third and fourth BcsB protomers in the c-di-GMP-saturated macrocomplex (Fig. 2b), and between the fourth and the fifth copy in the context of a dinucleotide-free synthase (Fig. 2c). Each BcsF copy features a single transmembrane helix, which upon exit from the inner membrane kinks into an amphipathic helical extension and a cytosolic C-terminal tail engaged in interactions with a BcsE N-terminal domain as described below.

**Fig. 2.**
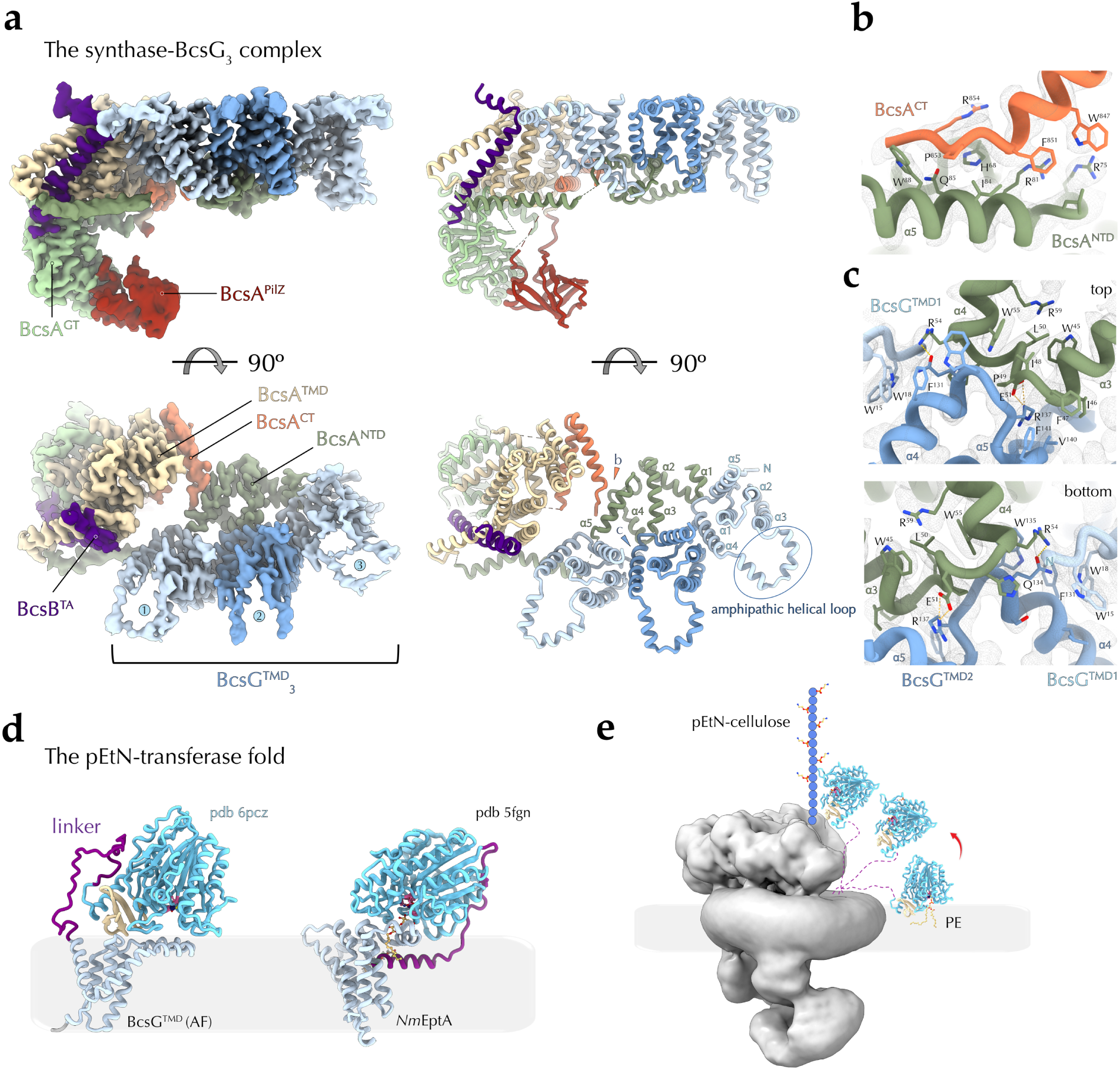
Cryo-EM structure of the synthase−pEtN-transferase complex. **a** Different views of a locally refined cryo-EM structure of the c-di-GMP-free BcsA-BcsB^TA^-BcsG_3_ assembly with corresponding electron densities (left) and cartoon representation (right). **b-c** Zoom-ins on the specific protein-protein interfaces with key residues shown as sticks and the electron density as a mesh. **d** Composite predicted structure of full-length BcsG (catalytic domain: X-ray structure of the *E. coli* BcsG^CTD^; NTD and linker, AlphaFold) and crystal structure of the lipid A pEtN-transferase from *Neisseria meningitidis* EptA. The flexible interdomain linkers are colored in purple. **e** Model for independent function of the three BcsG copies for substrate extraction and cellulose modification.

Using bacterial two-hybrid functional complementation (BACTH) assays we demonstrated previously that the *E. coli* type-specific N-terminal domain of the synthase interacts specifically with the BcsG enzyme *in cellulo*^11^. In addition, BcsG has been shown to directly affect BcsA integrity in the membrane^16,21^ and in some strains to be essential for cellulose secretion^11,21^. Indeed, using local refinement to resolve the more dynamically associated pEtN-transferase (Fig. 2 and Supplementary Fig. 3), we show here that BcsA^NTD^ adopts an amphipathic fold and recruits three copies of the BcsG pEtN-transferase whose transmembrane N-terminal domains are stabilized between BcsA^TMD^ and the sixth BcsB protomer of the crown, whereas the C-terminal catalytic BcsG modules remain unresolved in the structures. The BcsA N-terminus folds into a W-shaped series of α−helices whose connecting loops coordinate the BcsG protomers via conserved amino acid motifs in an otherwise weakly conserved primary structure (Fig. 2a-c). Each of the BcsG^NTD^ folds into 5 transmembrane helices (TMα1-5), which anchor the protein in the inner membrane and via the TMα4−TMα5 connecting loop interact with BcsA^NTD^, which in turn is stabilized by an α-helical amphipathic hairpin formed by the BcsA C-terminus (Fig. 2a-c). The BcsG TMα3−TMα4 linker region, on the other hand, folds into a short amphipathic α-helical loop at the periplasmic membrane interface, whereas TMα5 is predicted to extend into a 48 residue-long flexible linker^22^, followed by crystallographically characterized^21,23^ but here unresolved C-terminal pEtN-transferase domain (Fig. 2a and 2d). Based on homology with other alkaline phosphatases, such as the *Neisseria meningitidis* lipid A pEtN-transferase EptA (*Nm*EptA)^24^, the amphipathic helical loop and the extended interdomain linker likely assist the C-terminal catalytic domain in substrate-extraction by interactions with the polar headgroups of periplasm-facing phospholipids (phosphatidyl-ethanolamine (PE) in the case of BcsG) and allow for significant conformational flexibility in substrate delivery to the target acceptor. We show here that in BcsG the amphipathic helical loops point outwards relative to the crown lumen and proposed polysaccharide extrusion path, suggesting major conformational ‘gymnastics’ of the catalytic C-terminal domains for pEtN extraction and transfer onto the nascent cellulosic polymer (Fig. 2e). Such movements are likely made possible by the abovementioned interdomain linker and could explain the lack of resolved BcsG^CTD^-corresponding regions in the averaged electron density maps.

Interestingly, the presence of three BcsG copies is in contrast with a previous assignment of densities from a low-resolution cryo-EM map of the macrocomplex to a dimeric BcsG enzyme^16^ and most of the reported mechanistic studies on active pEtN-transferases − including on the C-terminal periplasmic module of BcsG − present no substrate- or product-determined prerequisite for catalytic domain oligomerization^21,23–29^. This suggests that the three BcsG copies visualized here likely act independently from each other to dynamically sample the membrane for, extract and transfer pEtN moieties from inner-membrane PE onto the nascent polysaccharide (Fig. 2e). Importantly, while this manuscript was under preparation a separate preprint reported independently the recruitment of trimeric BcsG via BcsA^NTD^, based on lower-resolution cryo-EM data, subcomplex purification and AlphaFold modeling^30^. Together, these results further validate the experimental structural data presented here and the two studies integrate and redress the structure-function model of pEtN-transferase association and function.

### The BcsF:BcsE interactions for cytosolic complex recruitment

We previously demonstrated that the cellulose secretion enhancer BcsE can form equimolar BcsE_2_R_2_Q_2_ complexes with the essential-for-secretion BcsRQ tandem in solution, that BcsE is sequestered by BcsF to the membrane and that BcsE’s N-terminal domain is necessary for stable cytosolic complex association with the synthase macrocomplex^14^. Nevertheless, how BcsE and BcsF interact, what structures they adopt in the secretory assembly and even their actual membrane-bound stoichiometries have remained unresolved^15,16^.

Here we show BcsE and BcsF interact in an asymmetric and heterotetrameric BcsE_2_F_2_ complex (Fig. 1b-c and Fig. 3a). In particular, BcsF adopts an X-shaped dimeric conformation within the inner membrane, stabilized by a hydrophobic N-proximal transmembrane interface burying 595 Å^2^ of surface area with free energy gain of −14.1 kcal/mol (Fig. 3b). At the C-termini, each BcsF protomer recruits a BcsE partner copy via cytosolic β-sheet complementation interactions with the central 9-stranded β-sheet of the interacting BcsE^NTD^ (Fig. 3b). The BcsF C-terminal tail threads along a shallow hydrophobic patch onto BcsE’s degenerate NTPase* domain and provides an overall charged solvent-exposed surface for the assembly (∼ 892 Å buried interface with free energy gain of −13.9 kcal/mol) (Fig. 3c). Consistent with the observed complex, BcsF truncations before or after P^43^ preceding the C-terminal cytosolic tail lead to incomplete Bcs macrocomplex assembly and corroborate the requirement for stable BcsF-BcsE^NTD^ interaction for vestibule complex recruitment (Fig. 3c). Importantly, the observed β-sheet complementation mechanism for BcsF-driven BcsE recruitment and BcsE^NTD^ dimerization (see below) is likely conserved across enterobacteria as shown in ColabFold and AlphaFold 3-predicted models of a consensus BcsE_2_F_2_ complex derived from representative homologs across the enterobacterial clade (Supplementary Fig. 4).

**Fig. 3.**
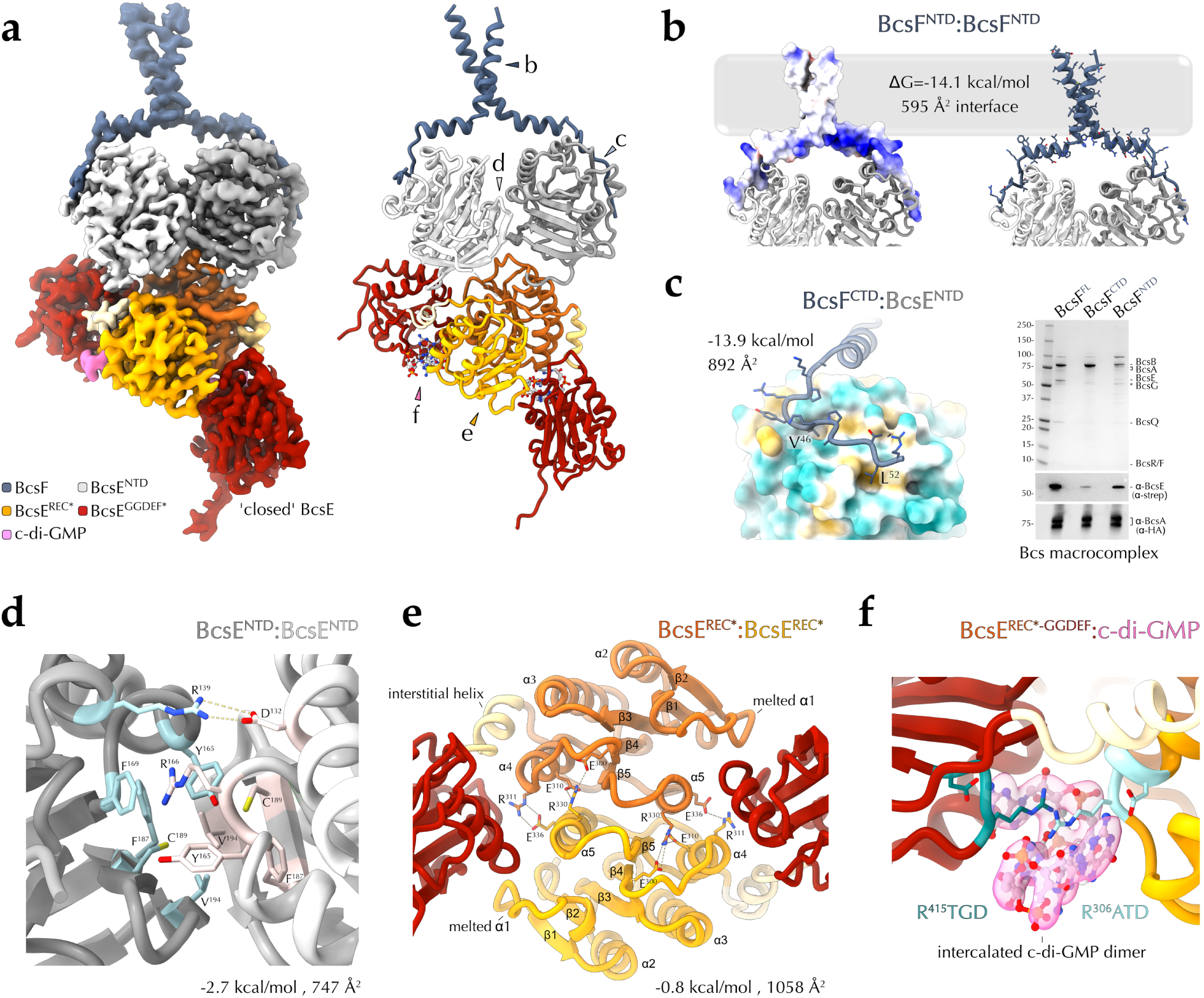
BcsF-dependent BcsE recruitment and regulatory complex conformation in the c-di-GMP-saturated state. **a** Locally refined cryo-EM structure of the BcsE_2_F_2_ assembly from the c-di-GMP-saturated synthase macrocomplex shown as electron density and in cartoon. **b** BcsF dimerization shown in as electrostatic potential-colored surface (left) and in cartoon and sticks (right). **c)** BcsE-BcsF interactions. Left, BcsE^NTD^ is shown as a hydrophobicity-colored surface, BcsF residues − including the hydrophobic plug residues V^46^ and L^52^ − are shown as sticks. Right, recombinant expression and purification of the Bcs macrocomplex with various BcsF variants (Bcs^His^RQA^HA-FLAG^B + Bcs^strep^EF*G). Protein-specific bands are identified as previously^11,15^. BcsE and BcsA-specific signals are further detected by western blotting with epitope tag-specific antibodies in the bottom. **d)** The BcsE^NTD^ dimerization interface. **e)** The BcsE^REC*^ dimerization interface. **f)** The c-di-GMP-binding dual I-site pocket in ‘closed’ BcsR. All interface parameters were calculated with PISA^51^.

The cryo-EM structures presented here are consistent with the previously tripartite architecture of BcsE, comprising a degenerate trio of an NTPase*, REC* and GGDEF* domains (Fig. 1a). Nevertheless, rather than engaging in head-to-tail interactions as proposed previously based on indirect BACTH interaction assays^14,15^, the two BcsE^NTD^ modules pack against each other in a head-to-head fashion, stabilized primarily by hydrophobic and π−stacking interactions in the center and by the peripheral BcsF C-terminal tails at the periphery (747 Å buried with free energy gain of −2.7 kcal/mol at the BcsE^NTD^ dimer interface) (Fig. 3d).

The REC*-GGDEF* domain tandem interacts with BcsQ via an extended C-terminal tail trailing along the BcsQ surface as observed in crystallographic snapshots previously^15^. However the REC* domains, which are not in contact in the crystallized states, engage in a head-to-head dimerization interactions mediated by a α4-β5-α5 interface (Fig. 3e), as observed as a canonical REC domain dimerization interface in many phosphorylation competent response regulators^31,32^. An intercalated c-di-GMP dimer is found at each *cis*-interdomain interface of a ‘closed’ BcsE^15^, stabilized by a composite R^306^ATD-R^415^TGD I-site tandem contributed by the corresponding REC* and GGDEF* domain, respectively (see below) (Fig. 1a and Fig. 3f). Finally, the two GGDEF* domains adopt different orientations relative to the to the apical BcsR_2_Q_2_ tandem consistent with the overall macrocomplex asymmetry. In the c-di-GMP-saturated macrocomplex, one BcsE^GGDEF*^ copy adopts an overall interaction interface consistent with the previously reported crystallized states and contacts BcsA^GT^ via its REC* module. The second BcsE^GGDEF*^ positions above the BcsQ dimer interface and is further stabilized by the β-strand connecting loops at the bottom of the BcsA^PilZ^ domain barrel (Fig. 3a). In the macrocomplex featuring a c-di-GMP-free synthase, the relative orientation of the REC* and GGDEF* BcsE modules are yet different and discussed below.

### The activating synthase : BcsRQ interactions

We previously showed that upon co-expression BcsR and BcsQ stabilize each other via the formation of a heterotetrameric BcsR_2_Q_2_ complex^14^, with essential roles in Bcs system positioning, assembly, stability and function^11,15,33^. Using X-ray crystallography and cryo-EM, we positioned the latter at the apical densities of the cytosolic vestibule around the synthase’s PilZ domain, however, the limited electron density map resolution prevented us from deciphering the specific protein-protein interactions and their roles in cellulose secretion^15^. Here we locally refined the structure of the crown-less Bcs macrocomplex complex to an average resolution of 2.85 Å, visualizing all interaction interfaces and coordinated nucleotide co-factors. An assymetric BcsR_2_Q_2_ complex is recruited to the membrane via BcsE’s C-terminal elongated tails, where both BcsQ copies interact with the synthase’s PilZ module (Fig. 4a-b) and adopt the nucleotide-driven sandwich dimer conformation characteristic for the SIMIBI (<u>si</u>gnal recognition particle, <u>Mi</u>nD, and <u>Bi</u>oD) family of protein-sorting NTPases to which BcsQ belongs^15^. Consistent with the previously reported crystal structures of the BcsR_2_Q_2_ complex^15^, the BcsR copies stabilize the ATP-bound BcsQ apical dimer via their V-shaped C-terminal tandem of α−helices (αC_1_ and αC_2_). Importantly, whereas one of the BcsR protomers is solvent-exposed and features an unstructured N-terminal domain, the other BcsR copy also interacts with the back of the BcsA catalytic module (Fig. 4a-c). Contrary to the crystallized states where BcsR^NTD^ adopts a β-hairpin conformation, here residues D^21^-S^30^ fold into an N-proximal α−helix (αN) that U-turns into an extended linker before adopting the V-shaped C-terminal domain onto the BcsQ dimer interface (Fig. 4c). The resulting N-terminal hairpin nestles into a hydrophobic BcsA^GT^ pocket via a L^25^-F^29^-L^31^-I^34^ plug at the tip and a I^22^:Y^36^ stabilizing interaction at the base. The strictly conserved D^21^ positions between R^367^ from the BcsA^GT^ domain and R^792^ in the middle of the C-proximal ‘hinge’ that enables PilZ rotation upon BcsA:c-di-GMP complexation (Fig. 4c). Overall, the BcsR:BcsA interaction interface buries 1059 Å and contributes a free energy gain of −5.8 kcal/mol (Fig. 4b-c). The BcsA^PilZ^ domain orientation is further stabilized by interactions between the β4-β5 connecting loop of the PilZ barrel and N-proximal residues from BcsR-αC_1_, as well as by an extensive interface with the underlying BcsQ protomer (682 Å buried with free energy gain of −4.1 kcal/mol) (Fig. 4b-c). On the other side of the β-barrel, an intercalated c-di-GMP dimer is found coordinated between the arginines from the canonical R^696^RxxR motif in the N-proximal PilZ domain linker, the active site gating loop is unstructured and the active site is substrate accessible (Fig. 4d). Together, the BcsEF-stabilized BcsRQA complex appears to lock the synthase into a catalytically competent state, which is consistent with previous *in vitro* activity data demonstrating dramatic synthase activation in the presence of excess cytosolic vestibule components, with stimulatory effects observed even in the absence of c-di-GMP^16^. Consistent with the observed BcsR:BcsA interactions, plasmid-based overexpression of BcsR leads to overproduction of matrix pEtN-cellulose (Fig. 4b), whereas mutations in the N-terminal domain, which do not affect BcsRQ complex formation *per se*, led to severe or complete loss of pEtN-cellulose secretion (Fig. 4e).

**Fig. 4.**
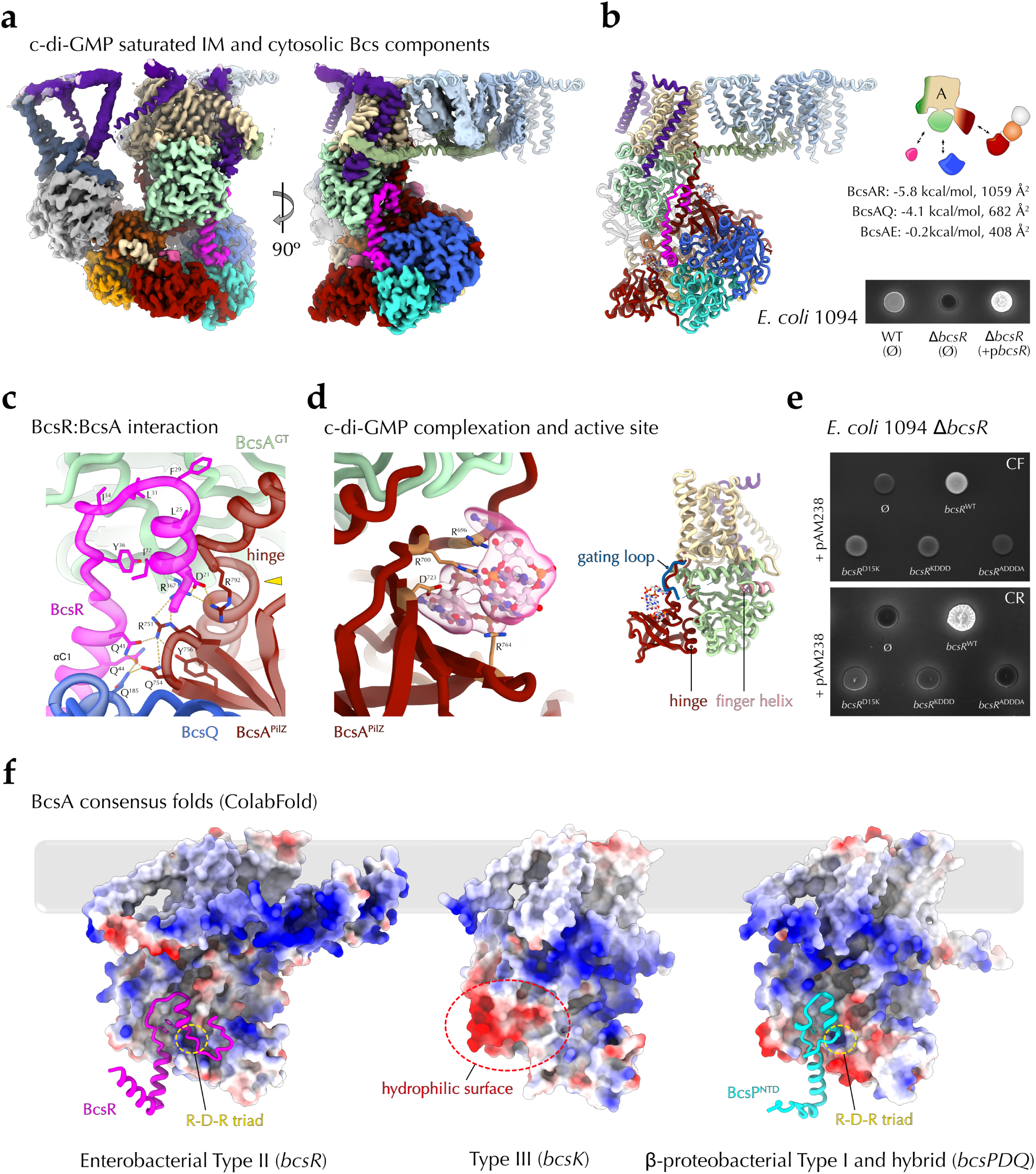
The c-di-GMP-bound synthase macrocomplex. **a** Locally refined cryo-EM map and fitted structure of the ‘crown’-less c-di-GMP-saturated synthase macrocomplex in two different views. **b** Cartoon representation of the same assembly, summary of the BcsA interactions with the cytosolic vestibule partners and stimulatory effects of BcsR overexpression as detected by binding and UV-fluorescence of E. coli macrocolonies grown on Congo Red-supplemented plates. **c** A zoom-in on the BcsA-BcsR interface with key residues shown as sticks. The R-D-R triad is indicated with a yellow arrowhead. **d** Effects on cellulose secretion upon BcsR^NTD^ mutagenesis using plasmid-based complementation with various BcsR mutants. KDDA, D^21^K-L^25^D-F^29^D-L^31^D; ADDDA, D^21^A-L^25^D-F^29^D-L^31^D-Y^36^A. CR, Congo Red; CF, calcofluor. **e** c-di-GMP coordination, together with its corresponding electron density, and overall core synthase fold showing unstructured gating loop and an accessible active site.

The experimentally determined BcsR:BcsA interaction interface via a surface-exposed hydrophobic pocket at the back of the synthase’s GT and PilZ modules is likely conserved across the enterobacterial clade. Indeed, a ColabFold model of a consensus BcsRA complex based on protein sequences from representative cellulose-secreting enterobacteria demonstrates an overall conserved fold and interface residues, including a hydrophobic plug at the tip, stabilizing F:Y π−stacking interactions at the base of the BcsR^NTD^ hairpin, and a conserved R:D:R triad at the GT:BcsR:‘hinge’ interface (Fig. 4f). Interestingly, similar fold prediction based on a consensus BcsA sequence derived from homologs encoded by *bcsK*-containing Type III *bcs* clusters^4,10^ lacks a corresponding hydrophobic pocket despite an overall high conservation of the BcsA sequence and fold (Fig. 4f).

We recently showed that most β-Proteobacteria featuring *bcsD* in a Type I or hybrid *bcs* operon architecture, also encode proline-rich BcsP homologs^17^. Similarly to the proline-rich cellulose crystallinity factor BcsH/CcpAx from *Gluconacetobacter hansenii*, which determines the formation of a longitudinal BcsHD cytoskeletal scaffold (a.k.a. ‘cortical belt’) and the respective linear alignment of the synthase terminal complexes for cellulose secretion and crystalline ribbon formation^34,35^, β-proteobacterial proline-rich BcsP recruits BcsD into distinct cytoskeletal assemblies that are key to cellulose biogenesis and the mature biofilm architecture^17^. Interestingly, the N-terminal regions of BcsP homologs show homology to enterobacterial BcsR^10^ and, similarly to the latter, BcsP expression and/or stability appeared enhanced in the presence of co-expressed and interacting BcsQ^17^. We therefore retrieved multiple sequences of representative and co-occurring β-proteobacterial BcsA and BcsP homologs and modeled the consensus complex between the synthase and BcsP^NTD^. Indeed, the predicted structure confirms both the presence of a conserved hydrophobic pocket on the synthase and a BcsR-like hairpin-shaped plug for BcsP^NTD^, suggesting a common mechanism for synthase regulation among widespread Type I and Type II Bcs secretion systems (Fig. 4f).

### BcsA activation-by-proxy in non-saturating c-di-GMP concentrations

BcsE was originally defined as a GIL-, or GGDEF I-site like-, domain protein due to a conserved C-terminal domain sensing c-di-GMP via an RxxD (R^415^TGD in *E. coli* BcsE) motif similar to the product-sensing I-sites found on many catalytically active diguanylate cyclases involved in feedback inhibition or dinucleotide signal relay^32,36^. We demonstrated previously that the GIL domain is in fact a degenerate and conformationally dynamic REC*-GGDEF* domain tandem where the R^415^TGD sequence corresponds to the canonical I-site in an otherwise catalytically incompetent module^14^. Whereas this motif is absolutely necessary for dinucleotide complexation, the phosphorylation-incompetent REC* domain can undergo significant conformational rearrangements to contribute a second I-site motif (R^306^ATD) for an intercalated c-di-GMP dimer complexation^15^ (Fig. 1a). This corresponds to a relatively compact or ‘closed’ BcsE conformation observed in a BcsRQE^REC*-GGDEF*^ crystal structure reported previously^15^ and also consistent with the cryo-EM structure presented above. The dissociation constants for dimeric c-di-GMP complexation thus changes from the low micromolar (∼2.5 μM, for the contribution of the GGDEF* I-site alone) to the nanomolar range (∼140 nM, for dual I-site coordination)^15^ (Fig. 1a). The latter c-di-GMP-binding affinity is significantly higher than the affinity for activating c-di-GMP complexation by the BcsA synthase itself, previously reported in the low micromolar range and orders of magnitude higher than the global cytosolic c-di-GMP concentrations in the early stages of biofilm formation^3,12,37^.

This raises the question of whether and how c-di-GMP binding to the higher-affinity sensor BcsE could have stimulatory effects on synthase activity and cellulose biogenesis in non-saturating dinucleotide concentrations. One possible mechanism is that the molecular ‘breathing’ of the Bcs macrocomplex during the processive cycles of glucose polymerization could cause reiterative conformational changes in BcsE and the synthase, thus leading to diametric changes in their respective dinucleotide binding affinities and c-di-GMP recycling for reiterative synthase activation. Alternatively, the higher affinity c-di-GMP binding to BcsE, associated with the latter’s compact conformation within the multicomponent cytosolic vestibule could stabilize the synthase in a catalytically competent conformation regardless of direct dinucleotide complexation.

To gain mechanistic insights, we kept low micromolar concentration of c-di-GMP throughout the Bcs macrocomplex purification and prepared the cryogrids after a final fast concentration step to approximately 2:1 dinucleotide-to-Bcs macrocomplex ratio. As these conditions are close to the predetermined dissociation constants for dimeric c-di-GMP complexation to both the BcsA and the BcsE GGDEF* domain alone (i.e. consistent with ‘splayed’ BcsE without contributions of the secondary REC* domain I-site to nucleotide binding)^12,15^ but more than an order of magnitude higher than that for compact, tandem I-site-contributing BcsE^15^, we hypothesized that they could allow us to capture either a BcsE-saturated/BcsA non-saturated state or, inversely, a splayed, non-saturated BcsE accompanying a c-di-GMP-bound synthase.

About half of the structurally resolved particles featured the fully c-di-GMP-saturated state shown above, where all three c-di-GMP-sensing subunits (BcsA and BcsE_2_) are bound to an intercalated dinucleotide dimer in a preserved 2:1 dinucleotide-to-protein binding site ratio. Interestingly, the remaining particles showed a c-di-GMP-free BcsA synthase and a more extended vestibule conformation (Fig. 1c, Fig. 5a-c and Supplementary Fig. 1) where the central BcsE^REC*^ domains engage in a different dimerization interface mediated by the pairs of β1-β2 connecting loops (melted α1 relative to canonical response regulators) and the C-proximal α-helices (canonical α5) (Fig. 5d). Densities for the PilZ-proximal GGDEF* domain feature markedly lower resolution (Supplementary Fig. 5), however, the conformation for both BcsE^REC*-GGDEF*^ tandems is still consistent with the ‘closed’ BcsE state and intercalated c-di-GMP complexation (Fig. 5c and 5e). The overall BcsE fold features a more extended conformation along the NTPase*-REC* domain linkers, neither BcsE protomer contacts the cytosolic synthase modules and the X-shaped BcsF dimer is found shifted between the fourth and fifth BcsB protomer as opposed to the c-di-GMP-saturated complex shown above (Fig. 1c). Nevertheless, the BcsRQ tandem is retained as an apical complex and the synthase-proximal BcsQ and BcsR protomers engage in similarly extensive contacts with the catalytic and PilZ modules (Fig 5f). The latter is only partially rotated around the ‘hinge’ helix relative to the c-di-GMP bound state (12.8° rotation and 1.7 Å displacement) and the N-proximal PilZ domain linker is partially unstructured but remains far from gating loop-stabilizing interactions with the BcsA^GT^ core. Conversely, the gating loop remains unresolved and active site appears substrate-accessible, suggesting an overall preserved catalytically competent state in the assembled macrocomplex (Fig. 5g).

**Fig. 5.**
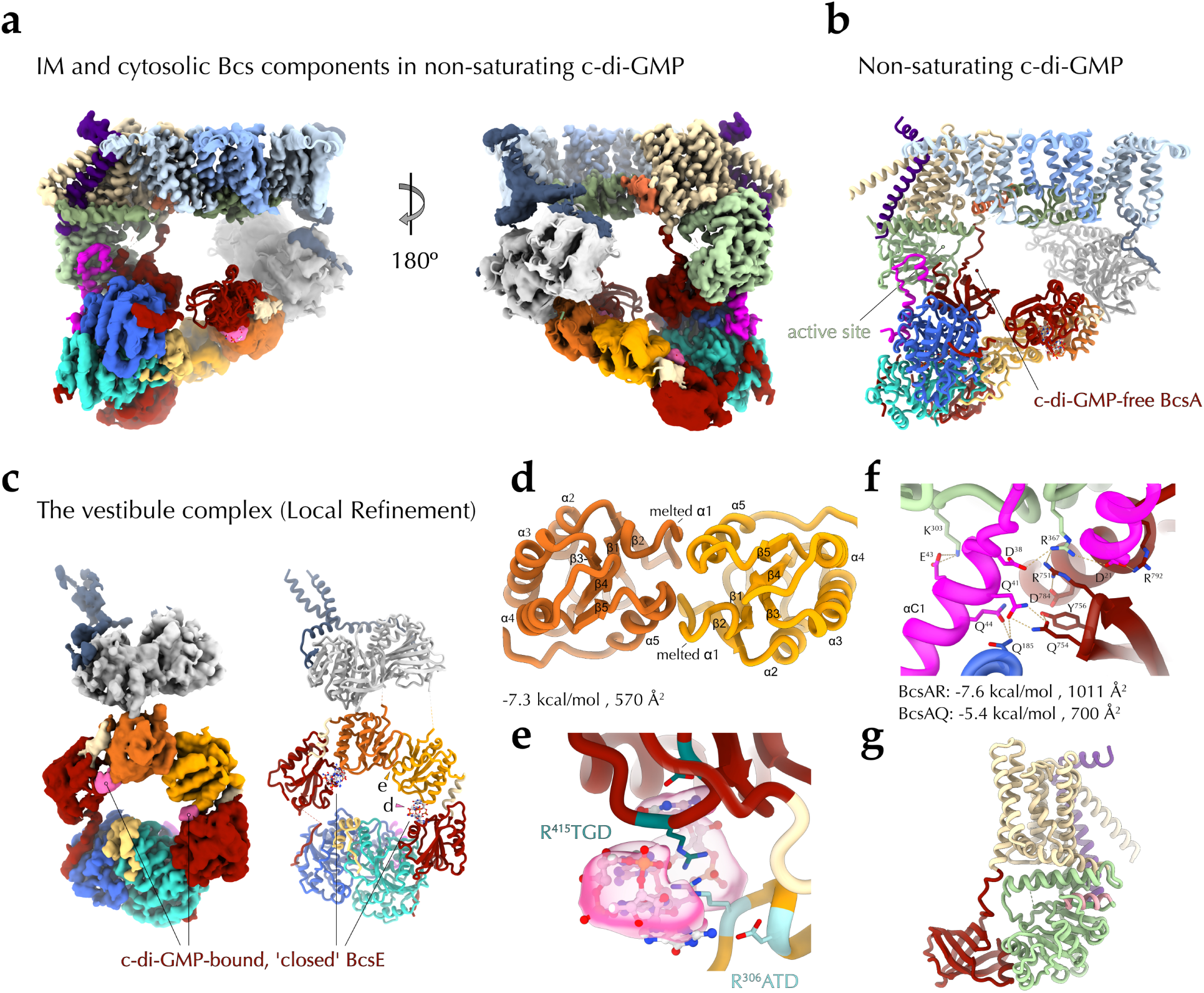
The synthase macrocomplex in limiting c-di-GMP. **a** Locally refined cryo-EM map and fitted structure of the ‘crown’-less synthase macrocomplex featuring a c-di-GMP-free synthase in two different views. **b** Cartoon representation of the same assembly. **c** The cryo-EM map and model of a locally refined BcsEFRQ assembly. **d** A zoom-in on c-di-GMP binding by a composite, dual I-site pocket in ‘closed’ BcsE. **e** REC* domain dimerization interface in the non-saturated macrocomplex. **f** A zoom-in on the BcsA:BcsR interface and summary of the synthase’s interactions with the cytosolic vestibule partners^51^. **g** Overall core synthase fold showing unstructured gating loop and an accessible active site.

Together, these data suggest that even lower, non-saturating c-di-GMP concentrations would allow dinucleotide binding to the nanomolar-affinity sensor BcsE via contributions of both its REC* and GGDEF* domain I-sites and would lead to sufficient BcsE compaction, assembly of the cytosolic vestibule and stabilization of the synthase modules in a BcsRQ-preactivated state. The specific BcsE REC* domain dimerization interface and overall vestibule conformation would also be likely influenced by the lateral diffusion and BcsF partner stabilization between the synthase-distal BcsB copies of the crown. Processive substrate addition and product release by BcsA would thus depend primarily on minute movements of its gating loop and finger helix - as observed *in crystallo* in saturating dinucleotide concentrations for the catalytic cycle of the *R. sphaeroides* BcsAB tandem - rather than on direct synthase-c-di-GMP complex formation. Overall, this is consistent with a model where the secondary c-di-GMP sensor BcsE serves as a proxy for dinucleotide-dependent regulation by effectively lowering the threshold for activating c-di-GMP concentrations and stabilizing the catalytically competent synthase state, rather than by circulating dinucleotide in and out of its PilZ-linker pocket (Fig. 6).

**Fig. 6.**
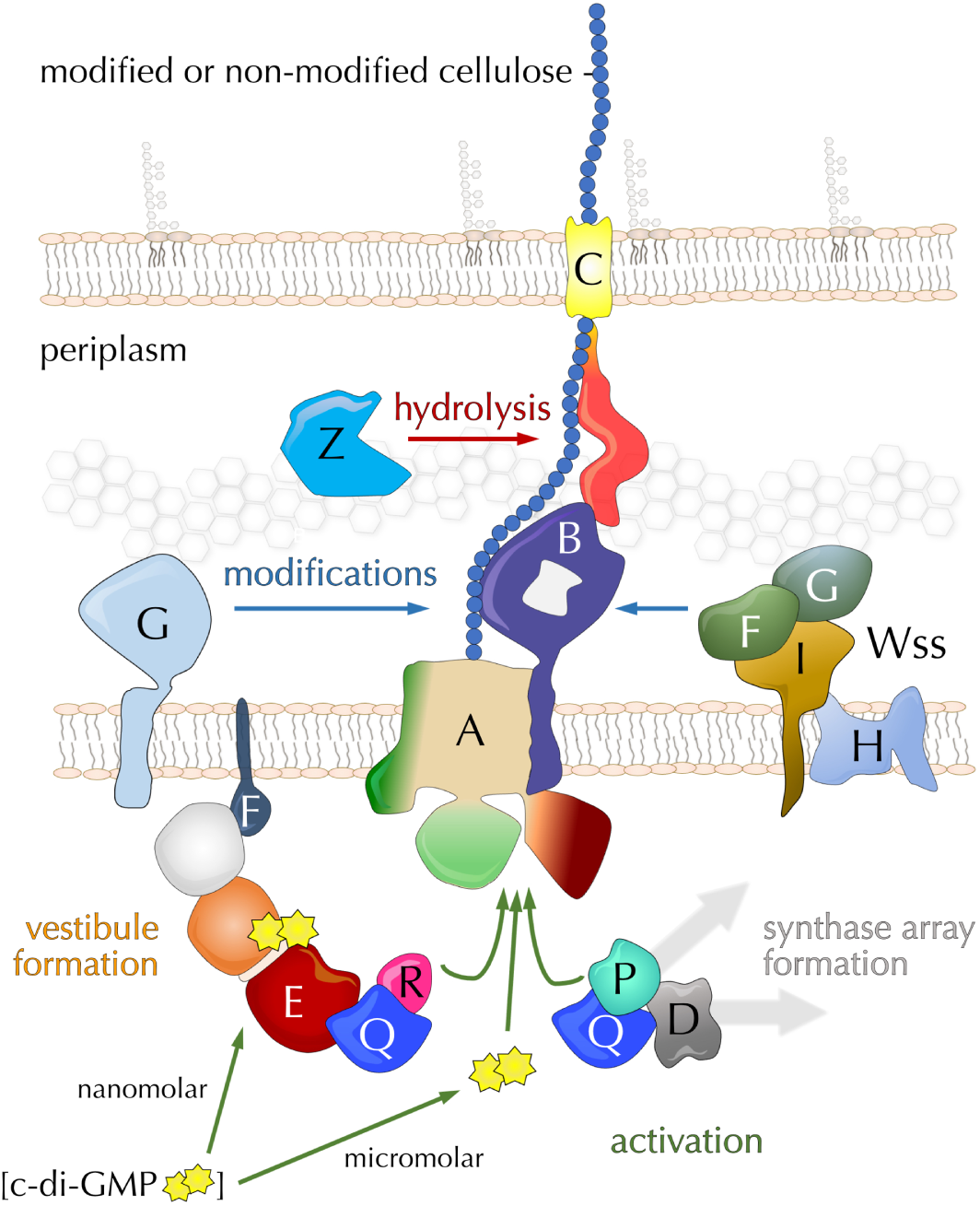
Synthase activation and polymer modifications in β- and γ−Proteobacteria. In addition to direct c-di-GMP complexation at micromolar dinucleotide concentrations, BcsA can be activated or stabilized in a catalytically competent conformation by a the high-affinity c-di-GMP sensing BcsEFRQ cytosolic vestibule complex or by macromolecular intracellular scaffolds. In the periplasm the polymer can undergo chemical modifications by the pEtN-transferase BcsG or by a multicomponent Wss cellulose acetylation complex. Finally, the polymer can undergo limited hydrolysis by the periplasmic endoglucanase BcsZ.

## Discussion

In many free-living and eukaryotic host-associated bacterial species, secreted cellulosic polymers represent key building components in the three-dimensional architecture of collaborative multicellular biofilms^3,6^. In *E. coli*, the phosphoethanolamine decoration of secreted cellulose influences not only the physicochemical properties of the polymer itself, but also favors higher-order, long-range fibrillation of the other major biofilm matrix component - amyloid curli - and thus provides for markedly increased biofilm cohesion and elasticity^38,39^. Importantly, the mature biofilm is a highly heterogeneous environment with stark gradients of oxygen, nutrients, moisture and/or shear stress, which leads to local stratification and/or compartmentalization of the quantity and type of secreted adherence factors and yields to a self-organized division of labor among subsets of cells engaged in extracellular matrix production, while others provide for cell proliferation and biofilm dispersal^3,40^.

In general, younger, nutrient-exposed biofilm layers are characterized by post-exponential growth metabolism, rod shape, enmeshed or no flagella, preserved proliferation and very low c-di-GMP levels (∼ 40-80 nM)^3^. Conversely, older, stationary phase strata cascade-activate specific subsets of diguanylate cyclases for gradual c-di-GMP increase, thus leading to non-dividing rounded cells embedded in a dense extracellular mesh of pEtN-cellulose and amyloid curli^3^. In intermediate layers, separate pockets or ridges of cells can activate specifically curli or pEtN-cellulose secretion, suggesting localized regulatory events that can selectively override the global c-di-GMP deficit^3^. Indeed, at least in some *E. coli* strains the Bcs macrocomplex has been shown to directly interact with a cellulose-specific diguanylate cyclase (DgcC/AdrA), which would dramatically increase the probability of c-di-GMP-BcsA encounter in comparison to dinucleotide diffusion from an overall depleted cytosolic pool^37^. In addition to the spatial sequestration of a pathway-specific diguanylate cyclase and in light of the structural data presented here, we propose that *E. coli* and related enterobacteria have evolved a highly cooperative nanomachine for efficient cellulose synthase activation and polymer modifications.

Consistent with our earlier but indirect BACTH results^11^, we reveal here that the *E.coli*-like BcsA synthases have evolved a specific N-terminal amphipathic domain that recruits three copies of the BcsG pEtN-transferase - an enzyme that is proposed to use inner membrane PE as a substrate for pEtN extraction and transfer via a S^278^-linked covalent intermediate, whose catalytic domain and enzymatic mechanism have been extensively characterized *in vitro* and *in vivo*^13,21,23^. It is important to note that whereas up to half of the glucose residues can be pEtN-modified in the processively secreted cellulose, PE is a small-headgroup zwitterionic phospholipid that is generally enriched in the inner, rather than the outer, leaflet of the inner membrane^41^. The recruitment of multiple BcsG copies could thus allow for efficient substrate mining for the extensive polymer modification, where the individual protomers are likely to act independent of each other. The membrane sampling and significant conformational changes, which would be required for pEtN-transfer onto a polymer processively extruded through the periplasmic BcsB crown, are likely enabled by the 48 amino-acid long interdomain linker that could potentially extend more than 10-15 nm in the periplasmic space and are consistent with the highly dynamic, large-scale structural transitions reported for other pEtN transferases such as the lipid A pEtN-transferase *Nm*EptA^24^.

In addition to recruitment of the BcsG complex, we further reveal the recruitment and interactions of the rest of the *E. coli*-characteristic Bcs subunits, which are either essential for (e.g. BcsRQ) or greatly affect cellulose biogenesis (BcsEF) *in vivo*^11^. We reveal that the hexameric BcsB crown, in addition to providing a likely polymer translocation pathway via luminally stacked CBDs in the crown’s lumen, also directly contributes to the stabilization of the trimeric pEtN-transferase complex and the BcsF transmembrane dimer via discreet interactions with its C-terminal tail anchors. We further demonstrate a cytosolic, BcsF-dependent β-sheet complementation mechanism for recruitment of the catalytically incompetent NTPase-like BcsE^NTD^, which itself leads to the stabilization of the entire BcsE_2_R_2_Q_2_ cytosolic vestibule around the catalytic and c-di-GMP sensing modules of the synthase. Although this vestibule is observed in two discreet c-di-GMP-bound conformations dependent on dinucleotide abundance and stabilizing BcsB-BcsF interactions, BcsA is BcsRQ-bound and, as a result, stabilized in a catalytically-competent conformation even in the absence of direct dinucleotide complexation.

Together, these data highlight two additional regulatory inputs for efficient synthase activation with widespread implications across enterobacteria and beyond. On one hand, the nanomolar-affinity, tandem I-site-presenting c-di-GMP sensor BcsE could effectively lower the threshold for activating c-di-GMP concentrations in the assembled Bcs macrocomplex (Fig. 6). Such activation by a separate c-di-GMP sensor is reminiscent of other widespread EPS secretion systems where activating c-di-GMP sensing is carried out either with the contributions of (e.g. in the poly-N-acetylglucosamine secretion system of *E. coli*) or fully by separate co-polymerase subunits (e.g. in the Pel or alginate secretion systems of *P. aeruginosa*) and in *E. coli* and other related bacteria could provide an important boost to cellulose secretion in the early stages and/or intermediate layers of biofilm development where cytosolic c-di-GMP is particularly low^3,5^. On the other, the observed PilZ-domain stabilizing BcsA-BcsR interactions are likely preserved in a wide range of BcsP-encoding Bsc secretion systems that do not necessarily feature a *bcsEFG* cluster but rather rely on BcsA-interacting BcsPDQ scaffolds for stabilization of the catalytically competent synthase state and enhanced polymer secretion^17^ (Fig. 6). Both the more widespread and idiosyncratic BcsA-regulatory mechanisms presented here can be harnessed for the selective targeting of a variety of cellulose secretion systems across free-living, pathogenic and symbiotic bacteria, as well as for the bioengineering of hybrid systems for the enhanced production of biotechnologically relevant polymers.

## Materials and Methods

The experiments were not randomized and the investigators were not blinded during experimental design, execution, or outcome assessment.

### Bacterial strains and plasmids

Oligonucleotides, construct design and bacterial strains are listed in Supplementary Tables S1 and S2. All plasmids for recombinant protein expression (see below) were propagated in and isolated from *E. coli* DH5a cells. Recombinant Bcs macrocomplex expression for structural studies was carried out in NiCo21(DE3) competent *E. coli* cells (New England Biolabs). Recombinant expression for assessment of BcsF roles in macrocomplex assembly was carried out in a T1 phage-resistant *Δbcs* BL21*(DE3) strain, featuring a deletion of both *bcs* operons (*bcsEFG* and *bcsRQAB*), as well as the corresponding interoperon region (see below). Phenotypic assays of colony morphology and calcofluor binding were carried out in the wild-type *E. coli* 1094 strain and the *E. coli* 1094 Δ*bcsR* strains transformed with variants of the low-copy pAM238 plasmid. All bacterial strains and plasmids used in this study are available upon request.

### Recombinant DNA techniques

DNA manipulations were carried out using standard protocols for polymerase chain reaction (PCR), molecular cloning, transformation, and DNA analysis. Procedures for cloning of *bcs*^HIS^*RQA*^HA-FLAG^*B* and *bcs*^Strep^*EFG* for co-expression from pACYCDuet1 and pRSFDuet1* are similar to those previously described. Briefly, the genomic region corresponding to *bcsRQA*^HA-FLAG^*B* was amplified using genomic DNA from the *E. coli* 1094 *bcsA^HA-FLAG^* strain as a template and a high-fidelity DNA polymerase (Phusion, New England Biolabs) with appropriate restriction sites introduced in the 5′ primer overhangs (sense/antisense PstI/NotI). In parallel, the pACYCDuet1 vector was also PCR-amplified to include the respective restriction sites for in-frame ligation under the pACYCDuet1 Promoter 1, including the incorporation of an N-terminal polyhistidine tag-coding sequence on *bcsR*. The genomic region corresponding to *bcsEFG* was PCR-amplified with appropriate restriction sites introduced in the 5′ primer overhangs (sense/antisense BamHI/NotI) and the pRFSDuet1 vector was amplified to introduce the respective restriction sites for in-frame ligation under the pRSFDuet1 Promoter 1 and to remove the polyhistidine tag-coding sequence (*). All PCR products were subsequently digested with the respective restriction enzyme pair (New England Biolabs), gel-purified, ligated using T4 DNA ligase (New England Biolabs), transformed into chemically competent DH5α cells, and plated on LB-agar plates containing an appropriate antibiotic (34 μg ml^−1^ chloramphenicol and 40 μg ml^−1^ kanamycin for the pACYCDuet-1 and the pRSFDuet-1 constructs, respectively). Single colonies were grown in 5 ml liquid LB medium at 37 °C overnight and the plasmid DNA was extracted using NucleoSpin® Plasmid preparation kit according to the manufacturer’s instructions (Macherey-Nagel). Positive clones were identified by restriction digestion and DNA sequencing. For introduction of a N-terminal STREP II tag-coding sequence in *bcsE*, the purified *bcsEFG-*pRSFDuet1* was inverse PCR-amplified with oligonucleotides including the epitope tag-coding sequence, the PCR product was gel-purified, 5’ phosphorylated using T4 polynucleotide kinase (New England Biolabs), ligated by the addition of T4 DNA ligase and transformed in *E. coli* DH5a cells for plasmid selection and amplification as above^15^.

### *E. coli* Δ*bcs* strain construction

The BL21*(DE3) Δ*bcs* mutant was generated using a modified protocol of a one-step inactivation procedure^42^. First, an FLP recognition target sites (FRT)-flanked kanamycin resistance (Km^R^) cassette was generated by PCR using the pKD4 plasmid as a template and a pair of oligonucleotides carrying 50-nucleotide extensions homologous to regions adjacent to the target *bcs* gene cluster. In parallel, BL21*(DE3) were transformed with the pKD46 plasmid and transformants were selected on LB-agar plates supplemented with 100 µg/ml ampicillin and grown at 30°C. Of these, a single colony was grown in liquid LB at 30°C, in the presence of ampicillin and 0.05% arabinose for induction of phage λ Red recombinase prior to chemically competent cells preparation. The PCR product was then transformed into the resulting BL21*(DE3) cells and transformants were selected on LB-agar plates supplemented with 40 µg/ml kanamycin and grown at 37°C, allowing for loss of the pKD46 helper plasmid. Replacement of the *bcs* gene cluster by the kanamycin-resistance cassette was confirmed by colony PCR. The resulting Δ*bcs*::Km^R^ strain was then transformed with the pCP20 helper plasmid, encoding Flp recombinase, and transformants were selected on ampicillin (100 µg/ml), then incubated for 24 h at 30°C to allow excision of the cassette by the expressed Flp recombinase. Plasmid pCP20 was then eliminated by growth at 37°C in the absence of antibiotics and the cells were verified for kanamycin and ampicillin sensitivity.

### Protein overexpression and purification

Overexpression of the Bcs macrocomplex was performed by coexpression of the pACYCDuet1-*bcs*^His^RQA^HA-FLAG^B and pRSFDuet1*-*bcs*^Strep^*EFG* constructs in chemically competent NiCo21(DE3) cells and plated on LB-agar plates with antibiotic concentrations reduced to two-thirds of the ones stated above. After overnight incubation of the plates at 37°C, multiple colonies of the transformed NiCo21(DE3) cells were picked and grown together at 37°C in antibiotics-supplemented terrific broth (TB) medium to optical density at 600 nanometers (OD_600_) of 0.8–1.2, upon which the cultures were transferred to 17°C and induced with 0.7 mM isopropyl-β-D-thiogalactopyranoside (IPTG, Neo Biotech) for 16h or overnight. Cells were pelleted by centrifugation (5000 g, 20 min, 4°C) and the pellets were resuspended in ice-cold buffer A containing 20 mM HEPES pH 8.0, 120 mM NaCl, 10 % glycerol, 5 mM MgCl_2_, 10 μM adenosine-5′-[(β,γ)-methyleno]triphosphate (AppCp, Jena Bioscience), 2 μM cyclic diguanylate (c-di-GMP, Sigma-Aldrich), 250 μM cellobiose, 0.5 mg ml^−1^ *Aspergillus niger* cellulase (Sigma-Aldrich), 100 μg ml^−1^ lysozyme, and 1 tablet per 50 ml complete EDTA-free protease inhibitors (Roche). The cells were subsequently disrupted using an Emulsiflex-C5 high-pressure homogenizer (Avestin) and the lysates were pre-cleared by a low-speed centrifugation step (10,000 g, 15 min, 4 °C). Membranes were pelleted by high-speed centrifugation using an SW 28 Ti or an SW 41 Ti Beckman rotor (26 500 rpm or 38 000 rpm, respectively, for 1h at 4 °C) and resuspended in solubilization buffer containing all buffer A components except for lysozyme and cellulase, as well as a mix of detergents at the following final concentrations: 0.6% w/v digitonin (Sigma-Aldrich), 0.35% w/v n-dodecyl-β-D-maltopyranoside (anagrade β-DDM, Anatrace), and 0.45% w/v lauryl maltose neopentyl glycol (LM-NPG, Anatrace). After incubation for 90 min at 22 °C and under mild agitation, the solubilized membrane fraction was cleared by a second high-speed centrifugation (50,000 g, 40 min, 4 °C). The supernatant was incubated with ANTI-FLAG® M2 affinity gel (100 μl resin per litre of induced culture, Sigma-Aldrich), under mild agitation at 4°C for 1 h. After gravity elution of the non-bound fraction, the resin was washed extensively (>30 column bed volumes) with affinity buffer containing 20 mM HEPES pH 8.0, 120 mM NaCl, 5 mM MgCl_2_, 10 μM AppCp, 4 μM c-di-GMP, 250 μM cellobiose and 0.01% w/v LM-NPG. The bound complexes were eluted using four column bed volumes of elution buffer (affinity buffer supplemented with 3X FLAG® peptide at 100 μg ml^−1^), concentrated on a 100 kDa cut-off Amicon® Ultra (MerckMillipore) centrifugal filter. Samples were analyzed by SDS-PAGE and western blots. For cryo-EM grid preparation, the Bcs macrocomplex was concentrated to ∼2-4 mg/ml, spotted on glow-discharged (ELMO, Cordouan Technologies) gold UltrAuFoil R 1.2/1.3 cryogrids, blotted, and plunge-frozen in liquid ethane using a Vitrobot Mark IV device (Thermo Fisher Scientific) at 4°C and 100% humidity.

### Cryoelectron microscopy and single-particle analysis (cryo-SPA)

Cryogrids were prescreened and optimized on the Elsa Talos Arctica transmission electron microscope (Thermo Fisher Scientific) at the European Institute of Chemistry and Biology (IECB, Bordeaux) operated at 200 kV and equipped with a Gatan K2 Summit direct electron detector. For structure resolution cryo-EM data was collected at the CM01 beamline at the European Synchrotron Radiation Facility (ESRF, Grenoble) on a Titan Krios transmission cryo-electron microscope, operated at 300 kV and equipped with a Gatan K3 direct electron detector and a Quantum LS imaging filter. 20 022 movies (two movies per grid hole, 50 frames per movie) were recorded in electron counting mode with a total electron dose per movie of 49.35 electrons/Å^2^, corrected pixel size of 0.839 Å/pixel, and defocus spread from −2.1 to −0.3 μm. The movies were motion-corrected using MotionCor2^43^ within the ESRF autoprocessing pipeline and the resulting micrographs were imported in CryoSPARC^44^ v4.4.1 for Patch-CTF correction and downstream processing. Particles were autopicked using the software’s Template Picker function and 2D templates as previously reported and, after extraction (box size 500 pixels, Fourier crop 200) and a round of 2D classification, a total of 1 359 795 particles with resolved structural features were selected for further processing. Ab-Initio Reconstruction and Heterogeneous Refinement among three classes yielded a model consistent with the previously reported Bcs macrocomplex structure integrating 834 077 or 61% of the preselected particles. The corresponding particles were re-extracted without downsampling and non-uniform refinement led to a 3D reconstruction featuring well-resolved crown densities and less-resolved inner-membrane and cytosolic regions. The hexameric BcsB periplasmic crown was locally refined after subtracting the inner-membrane and cytosolic densities from the particles dataset using the Particle Subtraction function. An inverse Particle Subtraction was also used to subtract the periplasmic densities from the initial particles dataset in order to retain only the inner-membrane and cytosolic regions. The latter subtracted particles were then subject to another round of Ab-Initio Reconstruction with three classes yielding two well-resolved classes corresponding to the c-di-GMP-bound and the c-di-GMP-free synthase, whereas a third class featured poorly resolved structural features. Each of the resulting classes were input as a search model for heterogeneous refinement (3D classification) of the full macrocomplex, yielding the two states for the assembled macrocomplex after another round of Ab-Initio modeling for each 3D class. The respective crown regions were subtracted again and regions of interest were further refined via Local Refinement jobs after map segmentation and mask generation within Chimera^45^. Additional map sharpening for density interpretation was performed using Deep EMhancer^46^ via the CryoSPARC interface. Atomic model building and refinements were performed iteratively using previously reported BcsB, BcsRQ and BcsE^REC*-GGDEF*^ structures^14,15^ and AlphaFold3^47^ or ColabFold^48^-generated models as inputs for manual building in Coot^49^ and automated real-space refinement in Phenix^50^. Interface analyses were carried out with the PISA server^51^. Details of the data collection and refinement statistics are listed in Supplementary Tables S3 and S4, and Supplementary Figs. 2, 3 and 5. Structure visualization was performed in ChimeraX^52^.

### SDS-PAGE and Western blot analyses

Protein fractions were analyzed by standard denaturing SDS-PAGE electrophoresis using 4 to 20% gradient mini-gels (Bio-Rad), InstantBlue Coomassie protein stain, and a Bio-Rad GelDoc Go Infinity imager. For Western blot analyses, SDS-PAGE–migrated proteins were directly transferred using a standard mini-gel transfer protocol, polyvinylidene difluoride membranes, and a Trans-blot Turbo transfer system (Bio-Rad). Blocking and antibody incubations were performed in the presence of 5% skim milk or bovine serum albumin (for STREP II tag detection) in TPBS (1X phosphate-buffered saline supplemented with 0.1% Tween-20 detergent); all washes between and after antibody incubations were performed with 1×TPBS buffer. Mouse anti-HA (hemagglutinin) (Thermo Fisher Scientific, #26183; dilution 1:1000) and mouse anti-STREP II (QIAGEN, #34850; dilution 1:1000) antibodies were used as primary antibodies; HRP–conjugated donkey anti-mouse antibody (Abcam, ab6728; dilution 1:10,000) was used as secondary antibody. Signals were visualized using the Clarity Western ECL substrate and a ChemiDoc imaging system (Bio-Rad).

### Consensus structures modeling

BcsA, BcsP, BcsE and BcsF protein sequences encoded by operons coding for BcsR-BcsE-BcsF (Type II *bcs* clusters, 20 representative sequences for each protein), BcsP-BcsD (Type I *bcs* clusters, 30 representative sequences for each protein) or BcsK (Type III *bcs* clusters, 121 representative sequences for BcsA) were identified with the help of webFlaGs^53^ and the STRING^54^ and NCBI Nucleotide databases and aligned separately using Clustal Omega^55^. The alignments were visualized in JalView^56^ and trimmed for non-conserved N- or C-terminal extensions and internal sequence gaps. The corresponding consensus sequences were then retrieved and the proteins or protein complexes were modeled using the AlphaFold^47^ or ColabFold^48^ web server and visualized in ChimeraX^52^.

### Calcofluor- and congo red-binding assays

To test for the functional effects of the BcsA-interacting BcsR region, chemically competent cells were prepared from an *E. coli* 1094 wild-type and Δ*bcsR* deletion strains. The latter were transformed with a low-copy-number plasmid (pAM-238) carrying none, wild-type or mutant *bcsR* genes and plated on LB agar plates (Miller) supplemented with 60 μg/ml streptomycin. Single colonies were inoculated in 3 ml LB-streptomycin medium left to grow overnight at 37°C with agitation. On the following morning, 4 μl of each culture was spotted onto low-salt LB agar plates (1.5 g/L NaCl) supplemented with streptomycin, 0.1 mM IPTG, and 0.02% calcofluor (fluorescent brightener 28; Sigma-Aldrich) or 25 μg/mL Congo Red (Sigma-Aldrich). The spots were allowed to air dry, and the plates were incubated at 30°C. After 24 h, the plates were photographed under brief illumination with long-wave UV light (365 nm) for calcofluor fluorescence and with a GelDoc Go imaging system (Bio-Rad) under trans-UVB illumination (UV tray and ethidium bromide mode) for pEtN-cellulose specific Congo Red fluorescence.

## Acknowledgments

We would like to thank current and former members of the Structural Biology of Biofilms group, and especially W. Abidi, A. Siroy and M. Decossas, for discussions, technical assistance and/or work peripheral to the project. This project has also benefited from and has directly contributed to the IECB cryo-EM platform and we are grateful to R. Linares and E. Kandiah for data collection assistance at the CM01 beamline at the ESRF (Grenoble, France) and to E. Cascales and J. M. Ghigo for plasmid and strain sharing. This research has received funding from the ERC Executive Agency under grant agreement 757507 BioMatrix-ERC-2017-StG (to P.V.K.) and the *Agence Nationale de Recherche* (ANR, France) under grant agreements CelluSec and T-ERC CoG 2024 (to P.V.K.). Finally, Itxaso Anso is supported by the Postdoctoral Program under the Order of June 20^th^ 2023 of the Ministry of Education (Basque Country, Spain), which regulates and coordinates new grants and grant renewals for the Advancement of Doctorate-level Investigators (*Programa Posdoctoral de Perfeccionamiento de Personal Investigador Doctor*).

## Author contributions

P.V.K. conceived the project. I.A., S.Z., T.S. and P.V.K. designed, performed, and optimized the experimental procedures. I.A., S.Z., T.S. and P.V.K. analyzed the data. I.A. and P.V.K. secured funding and wrote the paper.

## Data and materials availability

All data needed to evaluate the conclusions in the paper are present in the paper and/or the Supplementary Information. Refined structural models and electron density maps are deposited in the electron microscopy and protein databanks and will be released upon peer-reviewed publication.

## References

1. Moradali, M. F. & Rehm, B. H. A. Bacterial biopolymers: from pathogenesis to advanced materials. Nat Rev Microbiol 18, 195–210 (2020).

2. Hobley, L., Harkins, C., MacPhee, C. E. & Stanley-Wall, N. R. Giving structure to the biofilm matrix: an overview of individual strategies and emerging common themes. FEMS Microbiol Rev 39, 649–669 (2015).

3. Serra, D. O. & Hengge, R. Bacterial Multicellularity: The Biology of *Escherichia coli* Building Large-Scale Biofilm Communities. Annu. Rev. Microbiol. 75, 269–290 (2021).

4. Krasteva, P. V. Bacterial synthase-dependent exopolysaccharide secretion: a focus on cellulose. Current Opinion in Microbiology 79, 102476 (2024).

5. Krasteva, P. V. & Sondermann, H. Versatile modes of cellular regulation via cyclic dinucleotides. Nat Chem Biol 13, 350–359 (2017).

6. Abidi, W., Torres-Sánchez, L., Siroy, A. & Krasteva, P. V. Weaving of bacterial cellulose by the Bcs secretion systems. FEMS Microbiology Reviews 46, fuab051 (2022).

7. Purushotham, P., Ho, R. & Zimmer, J. Architecture of a catalytically active homotrimeric plant cellulose synthase complex. Science 369, 1089–1094 (2020).

8. Ross, P. et al. Regulation of cellulose synthesis in Acetobacter xylinum by cyclic diguanylic acid. Nature 325, 279–281 (1987).

9. Morgan, J. L. W., McNamara, J. T. & Zimmer, J. Mechanism of activation of bacterial cellulose synthase by cyclic di-GMP. Nat Struct Mol Biol 21, 489–496 (2014).

10. Römling, U. & Galperin, M. Y. Bacterial cellulose biosynthesis: diversity of operons, subunits, products, and functions. Trends in Microbiology 23, 545–557 (2015).

11. Krasteva, P. V. et al. Insights into the structure and assembly of a bacterial cellulose secretion system. Nat Commun 8, 2065 (2017).

12. Omadjela, O. et al. BcsA and BcsB form the catalytically active core of bacterial cellulose synthase sufficient for in vitro cellulose synthesis. Proc. Natl. Acad. Sci. U.S.A. 110, 17856– 17861 (2013).

13. Thongsomboon, W. et al. Phosphoethanolamine cellulose: A naturally produced chemically modified cellulose. Science 359, 334–338 (2018).

14. Zouhir, S., Abidi, W., Caleechurn, M. & Krasteva, P. V. Structure and Multitasking of the c-di-GMP-Sensing Cellulose Secretion Regulator BcsE. mBio 11, e01303–20 (2020).

15. Abidi, W., Zouhir, S., Caleechurn, M., Roche, S. & Krasteva, P. V. Architecture and regulation of an enterobacterial cellulose secretion system. Sci. Adv. 7, eabd8049 (2021).

16. Acheson, J. F., Ho, R., Goularte, N. F., Cegelski, L. & Zimmer, J. Molecular organization of the E. coli cellulose synthase macrocomplex. Nat Struct Mol Biol 28, 310–318 (2021).

17. Sana, T. G. et al. Structures and roles of BcsD and partner scaffold proteins in proteobacterial cellulose secretion. Curr Biol 34, 106–116.e6 (2024).

18. Morgan, J. L. W., Strumillo, J. & Zimmer, J. Crystallographic snapshot of cellulose synthesis and membrane translocation. Nature 493, 181–186 (2013).

19. Morgan, J. L. W. et al. Observing cellulose biosynthesis and membrane translocation in crystallo. Nature 531, 329–334 (2016).

20. Du, J., Vepachedu, V., Cho, S. H., Kumar, M. & Nixon, B. T. Structure of the Cellulose Synthase Complex of Gluconacetobacter hansenii at 23.4 Å Resolution. PLoS One 11, e0155886 (2016).

21. Sun, L. et al. Structural and Functional Characterization of the BcsG Subunit of the Cellulose Synthase in Salmonella typhimurium. Journal of Molecular Biology 430, 3170–3189 (2018).

22. Varadi, M. et al. AlphaFold Protein Structure Database: massively expanding the structural coverage of protein-sequence space with high-accuracy models. Nucleic Acids Research 50, D439–D444 (2022).

23. Anderson, A. C., Burnett, A. J. N., Hiscock, L., Maly, K. E. & Weadge, J. T. The Escherichia coli cellulose synthase subunit G (BcsG) is a Zn2+-dependent phosphoethanolamine transferase. Journal of Biological Chemistry 295, 6225–6235 (2020).

24. Anandan, A. et al. Structure of a lipid A phosphoethanolamine transferase suggests how conformational changes govern substrate binding. Proc. Natl. Acad. Sci. U.S.A. 114, 2218– 2223 (2017).

25. Lu, D. et al. Structure-based mechanism of lipoteichoic acid synthesis by *Staphylococcus aureus* LtaS. Proc. Natl. Acad. Sci. U.S.A. 106, 1584–1589 (2009).

26. Schirner, K., Marles-Wright, J., Lewis, R. J. & Errington, J. Distinct and essential morphogenic functions for wall- and lipo-teichoic acids in Bacillus subtilis. EMBO J 28, 830–842 (2009).

27. Campeotto, I. et al. Structural and Mechanistic Insight into the Listeria monocytogenes Two-enzyme Lipoteichoic Acid Synthesis System. Journal of Biological Chemistry 289, 28054– 28069 (2014).

28. Clairfeuille, T. et al. Structure of the essential inner membrane lipopolysaccharide–PbgA complex. Nature 584, 479–483 (2020).

29. Fan, J., Petersen, E. M., Hinds, T. R., Zheng, N. & Miller, S. I. Structure of an Inner Membrane Protein Required for PhoPQ-Regulated Increases in Outer Membrane Cardiolipin. mBio 11, e03277–19 (2020).

30. Verma, P., Ho, R., Chambers, S. A., Cegelski, L. & Zimmer, J. Molecular insights into phosphoethanolamine cellulose formation and secretion. Preprint at 10.1101/2024.04.04.588173 (2024).

31. Gao, R. & Stock, A. M. Biological Insights from Structures of Two-Component Proteins. Annu. Rev. Microbiol. 63, 133–154 (2009).

32. Krasteva, P. V., Giglio, K. M. & Sondermann, H. Sensing the messenger: The diverse ways that bacteria signal through c-di-GMP: The Ins and Outs of c-di-GMP Signaling. Protein Science 21, 929–948 (2012).

33. Le Quéré, B. & Ghigo, J.-M. BcsQ is an essential component of the *Escherichia coli* cellulose biosynthesis apparatus that localizes at the bacterial cell pole. Molecular Microbiology 72, 724–740 (2009).

34. Nicolas, W. J., Ghosal, D., Tocheva, E. I., Meyerowitz, E. M. & Jensen, G. J. Structure of the Bacterial Cellulose Ribbon and Its Assembly-Guiding Cytoskeleton by Electron Cryotomography. J Bacteriol 203, (2021).

35. Abidi, W. et al. Bacterial crystalline cellulose secretion via a supramolecular BcsHD scaffold. Sci. Adv. 8, eadd1170 (2022).

36. Fang, X. et al. GIL, a new c-di-GMP-binding protein domain involved in regulation of cellulose synthesis in enterobacteria: GIL, a new c-di-GMP-binding domain. Molecular Microbiology 93, 439–452 (2014).

37. Richter, A. M. et al. Local c-di-GMP Signaling in the Control of Synthesis of the E. coli Biofilm Exopolysaccharide pEtN-Cellulose. Journal of Molecular Biology 432, 4576–4595 (2020).

38. Hollenbeck, E. C. et al. Phosphoethanolamine cellulose enhances curli-mediated adhesion of uropathogenic *Escherichia coli* to bladder epithelial cells. Proc. Natl. Acad. Sci. U.S.A. 115, 10106–10111 (2018).

39. Tyrikos-Ergas, T. et al. Synthetic phosphoethanolamine-modified oligosaccharides reveal the importance of glycan length and substitution in biofilm-inspired assemblies. Nat Commun 13, 3954 (2022).

40. Jo, J., Price-Whelan, A. & Dietrich, L. E. P. Gradients and consequences of heterogeneity in biofilms. Nat Rev Microbiol 20, 593–607 (2022).

41. Bogdanov, M. et al. Phospholipid distribution in the cytoplasmic membrane of Gram-negative bacteria is highly asymmetric, dynamic, and cell shape-dependent. Sci. Adv. 6, eaaz6333 (2020).

42. Datsenko, K. A. & Wanner, B. L. One-step inactivation of chromosomal genes in *Escherichia coli* K-12 using PCR products. Proc. Natl. Acad. Sci. U.S.A. 97, 6640–6645 (2000).

43. Zheng, S. Q. et al. MotionCor2: anisotropic correction of beam-induced motion for improved cryo-electron microscopy. Nat Methods 14, 331–332 (2017).

44. Punjani, A., Rubinstein, J. L., Fleet, D. J. & Brubaker, M. A. cryoSPARC: algorithms for rapid unsupervised cryo-EM structure determination. Nat Methods 14, 290–296 (2017).

45. Pettersen, E. F. et al. UCSF Chimera : A visualization system for exploratory research and analysis. J. Comput. Chem. 25, 1605–1612 (2004).

46. Sanchez-Garcia, R. et al. DeepEMhancer: a deep learning solution for cryo-EM volume post-processing. Commun Biol 4, 874 (2021).

47. Abramson, J. et al. Accurate structure prediction of biomolecular interactions with AlphaFold 3. Nature (2024) doi:10.1038/s41586-024-07487-w.

48. Mirdita, M. et al. ColabFold: making protein folding accessible to all. Nat Methods 19, 679– 682 (2022).

49. Emsley, P., Lohkamp, B., Scott, W. G. & Cowtan, K. Features and development of *Coot*. Acta Crystallogr D Biol Crystallogr 66, 486–501 (2010).

50. Adams, P. D. et al. PHENIX : a comprehensive Python-based system for macromolecular structure solution. Acta Crystallogr D Biol Crystallogr 66, 213–221 (2010).

51. Krissinel, E. & Henrick, K. Inference of Macromolecular Assemblies from Crystalline State. Journal of Molecular Biology 372, 774–797 (2007).

52. Pettersen, E. F. et al. UCSF CHIMERAX : Structure visualization for researchers, educators, and developers. Protein Science 30, 70–82 (2021).

53. Saha, C. K., Sanches Pires, R., Brolin, H., Delannoy, M. & Atkinson, G. C. FlaGs and webFlaGs: discovering novel biology through the analysis of gene neighbourhood conservation. Bioinformatics 37, 1312–1314 (2021).

54. Szklarczyk, D. et al. The STRING database in 2023: protein-protein association networks and functional enrichment analyses for any sequenced genome of interest. Nucleic Acids Res 51, D638–D646 (2023).

55. Sievers, F. et al. Fast, scalable generation of high-quality protein multiple sequence alignments using Clustal Omega. Molecular Systems Biology 7, 539 (2011).

56. Waterhouse, A. M., Procter, J. B., Martin, D. M. A., Clamp, M. & Barton, G. J. Jalview Version 2—a multiple sequence alignment editor and analysis workbench. Bioinformatics 25, 1189–1191 (2009).

