## Supplementary Information for "Structural basis for synthase activation and cellulose modification in the *E. coli* Type II Bcs secretion system"

### for

This document includes:

Supplementary Fig. 1-5

Supplementary Tables S1-S4

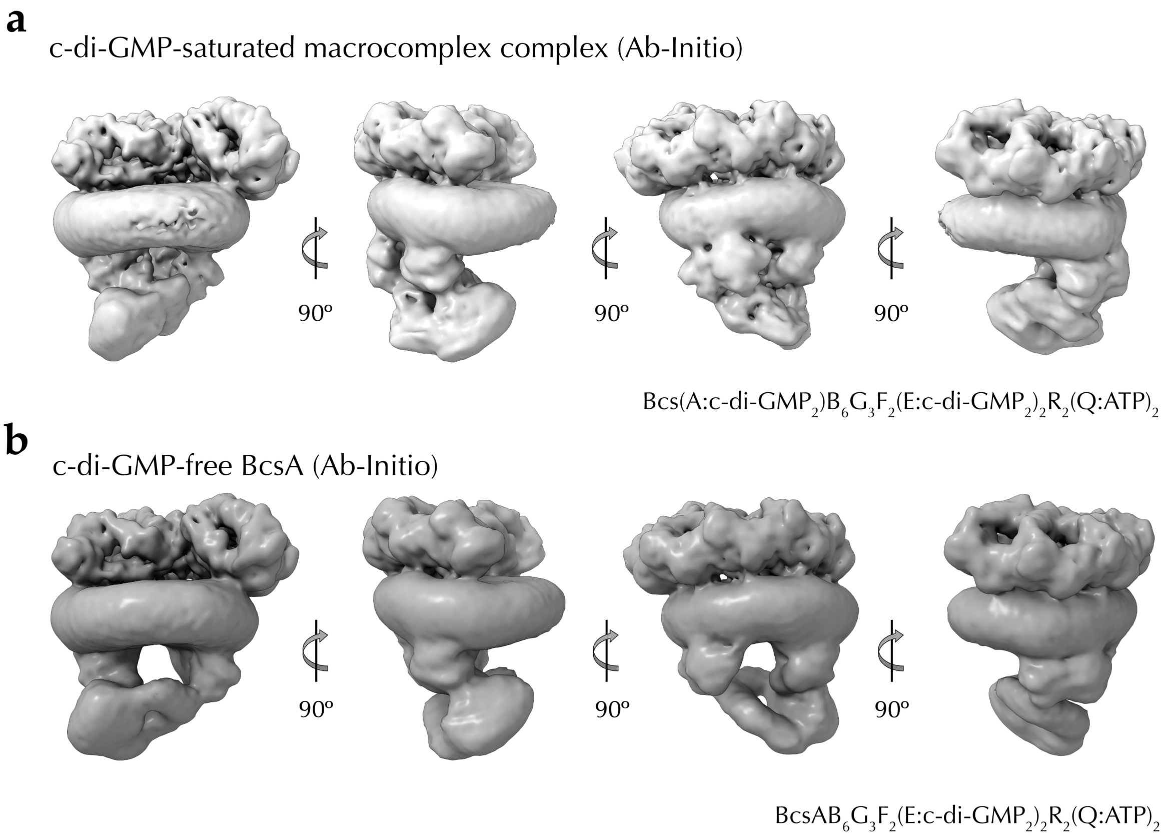


**Supplementary Fig. 1 Global architecture of the *E. coli* Bcs macrocomplex.** The two captured states, featuring a c-di-GMP-bound (**a**) or a c-di-GMP-free (**b**) BcsA synthase are shown as Ab-Initio electron density map reconstructions at ~7 Å resolution

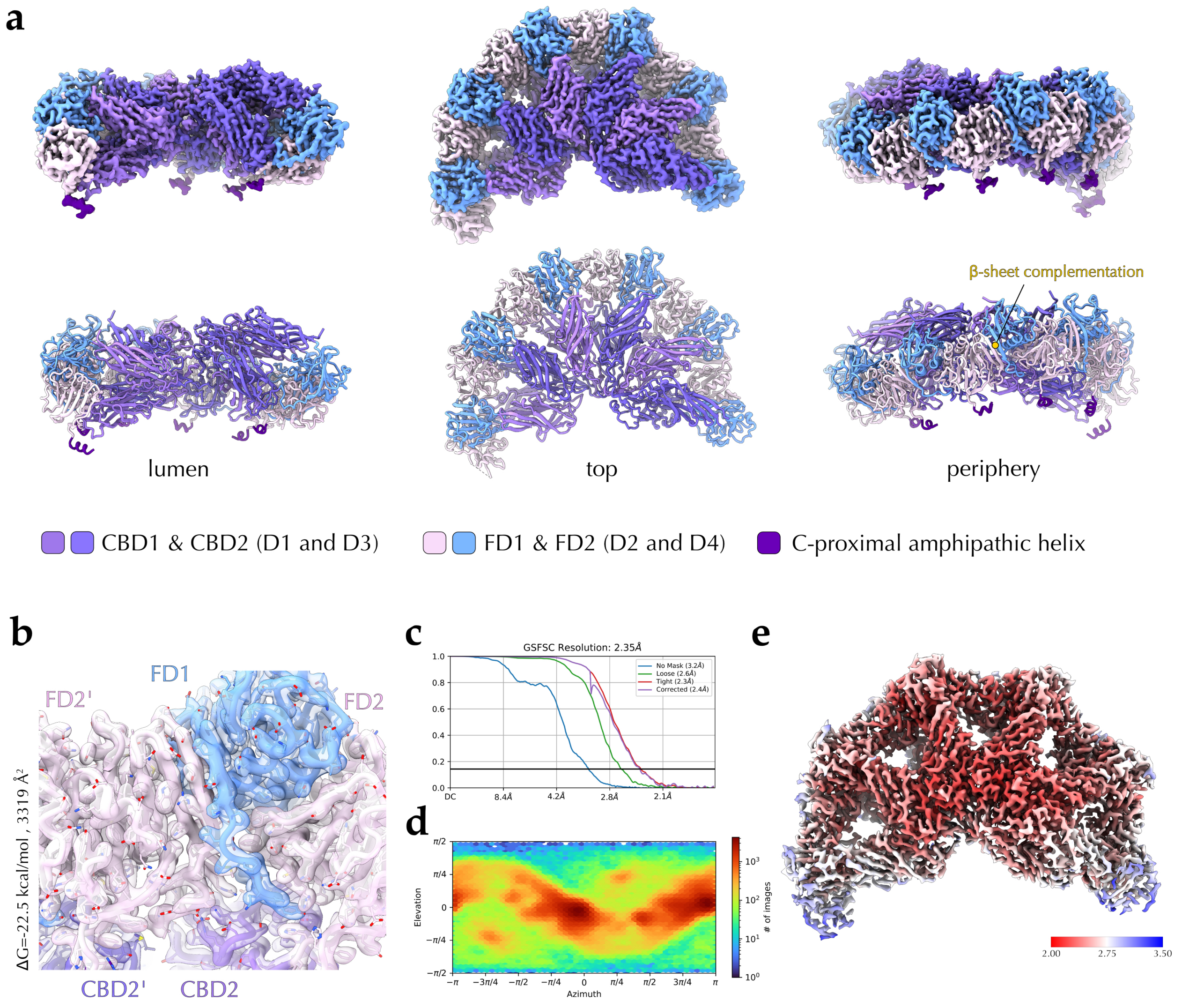


**Supplementary Fig. 2 Cryo-EM structure of the *E. coli* BcsB periplasmic crown. a** Cryo-EM density map and atomic model (in cartoon). D, domain; CBD, carbohydrate-binding domain; FD, flavodoxin-like domain. **b** a close-up of the model-in-map at the interprotomer interface. Free energy gain and buried surface area calculated by the PISA server. **c** Gold-standard Fourier shell correlation (GSFSC) curve used to determine the average density map resolution (CryoSPARC). **d** Viewing direction distribution across the dataset. **e** Surface mapping of the local resolution (2-3.50 Å in a red-blue gradient).

**a**

Non-saturated state (c-di-GMP-free synthase; Local Refinement)

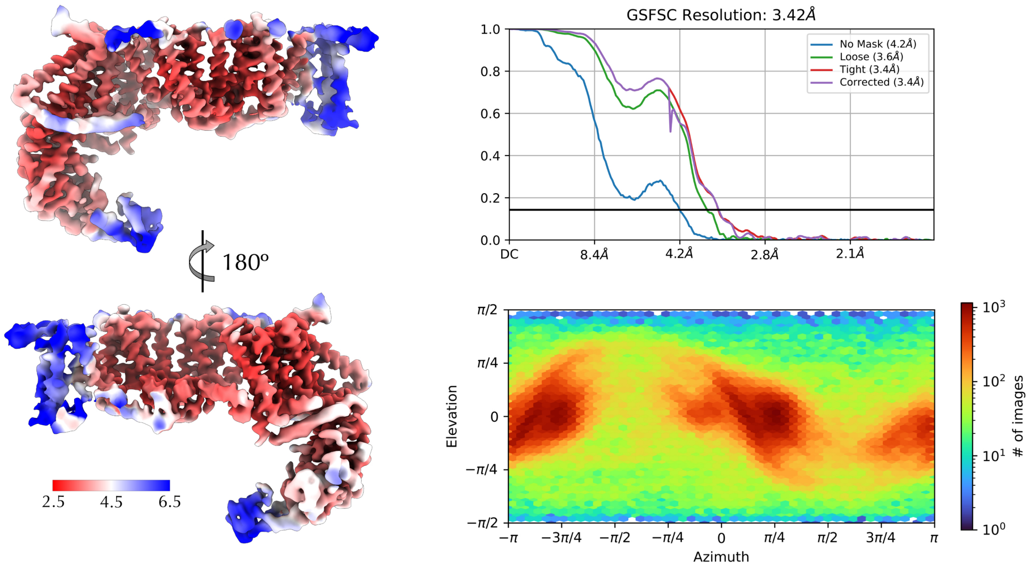


**b**

$BcsAG_3$ in the c-di-GMP-saturated Bcs macrocomplex (Local Refinement)

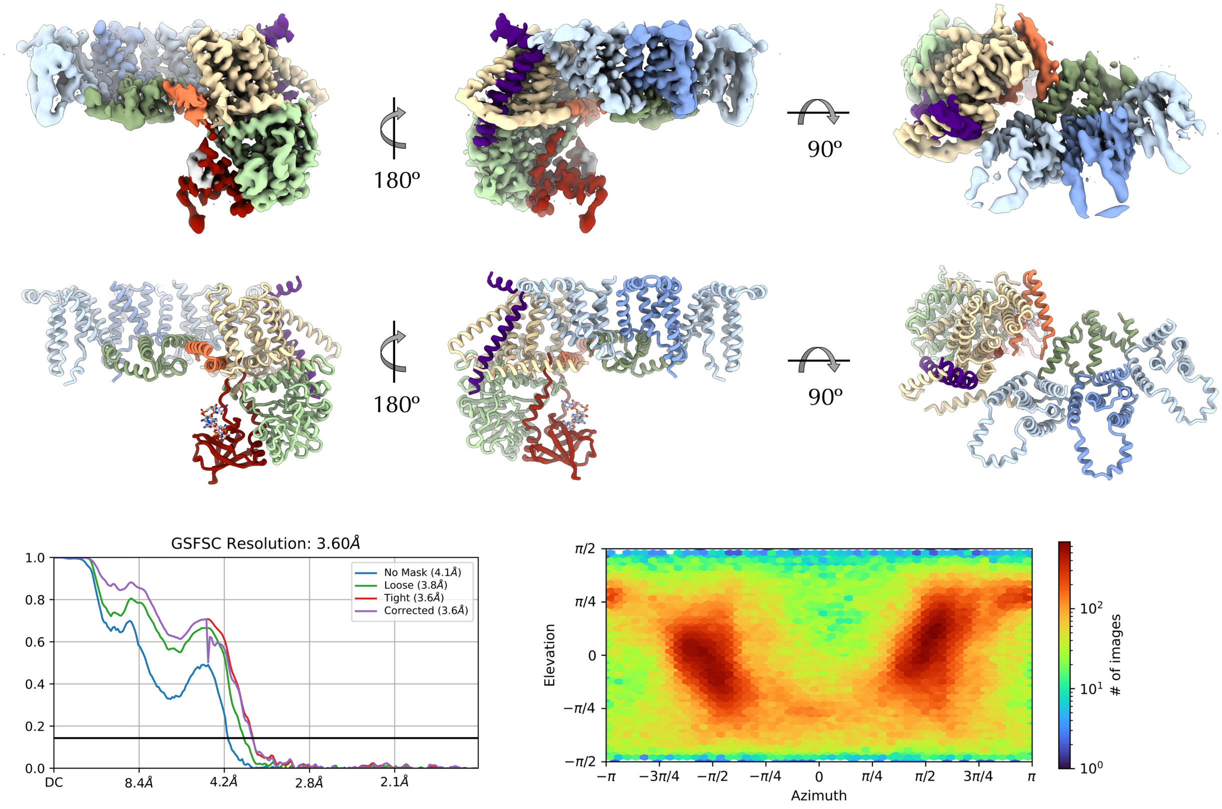


**Supplementary Fig. 3 Cryo-EM structures of the $BcsAG_3$ assemblies : supplementary information. a** Left, local resolution mapping as a red-blue color gradient onto the sharpened cryo-EM map of the c-di-GMP-free $BcsAG_3$ assembly. Right, the GSFSC curve and viewing direction distribution across the dataset. **b** Cryo-EM structure of the c-di-GMP-bound $BcsAG_3$ complex shown as a locally refined electron density map and an atomic model in cartoon. Bottom, GSFSC curve and viewing direction distribution.

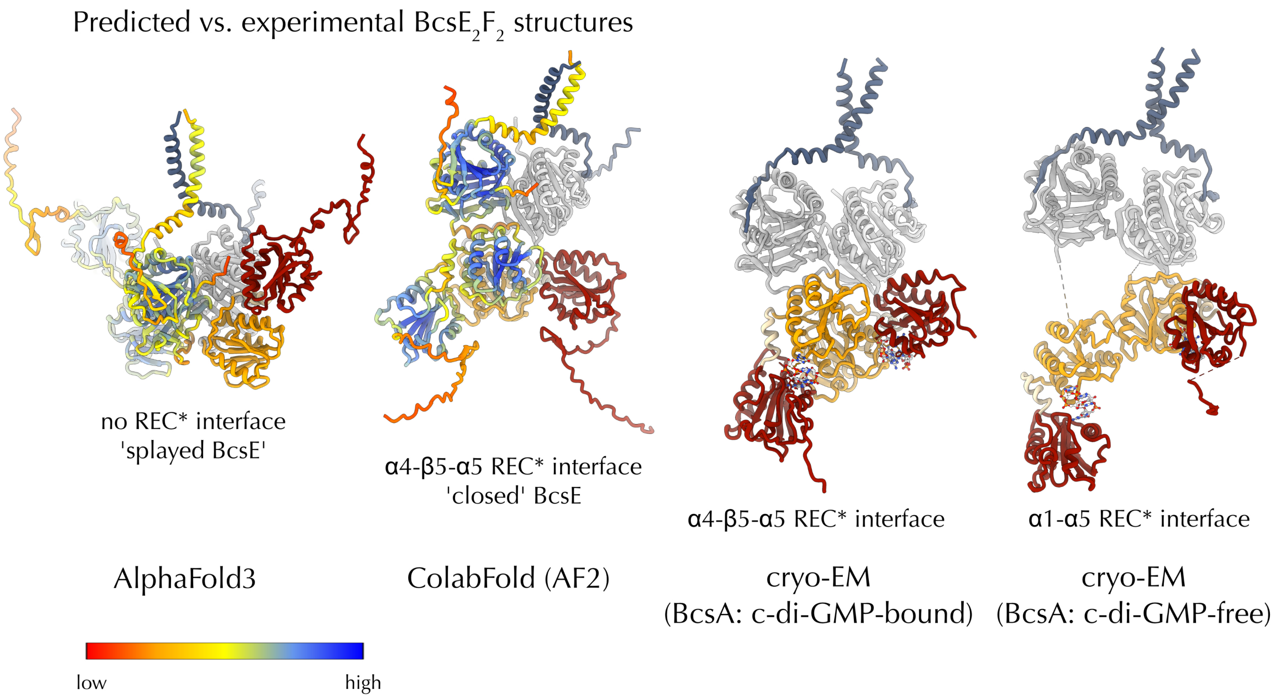


**Supplementary Fig. 4 Predicted vs. experimental structures of enterobacterial $E_2F_2$ regulatory complex.** From left to right: an AlphaFold 3-based model using consensus BcsE and BcsF sequences from representative cellulose-secreting enterobacteria; a ColabFold-predicted model using the same input sequences; cryo-EM structure of BcsEF in the c-di-GMP-saturated Bcs macrocomplex from *E. coli* (this study); cryo-EM structure of the same assembly in complex with a c-di-GMP-free BcsE (this study). Whereas head-to-head $BcsE^{NTD}$ dimerization and peripheral β-sheet complementation by BcsF are present in both predicted models, the experimental structures reveal additional conformational space for the BcsE cellulose secretion enhancer.

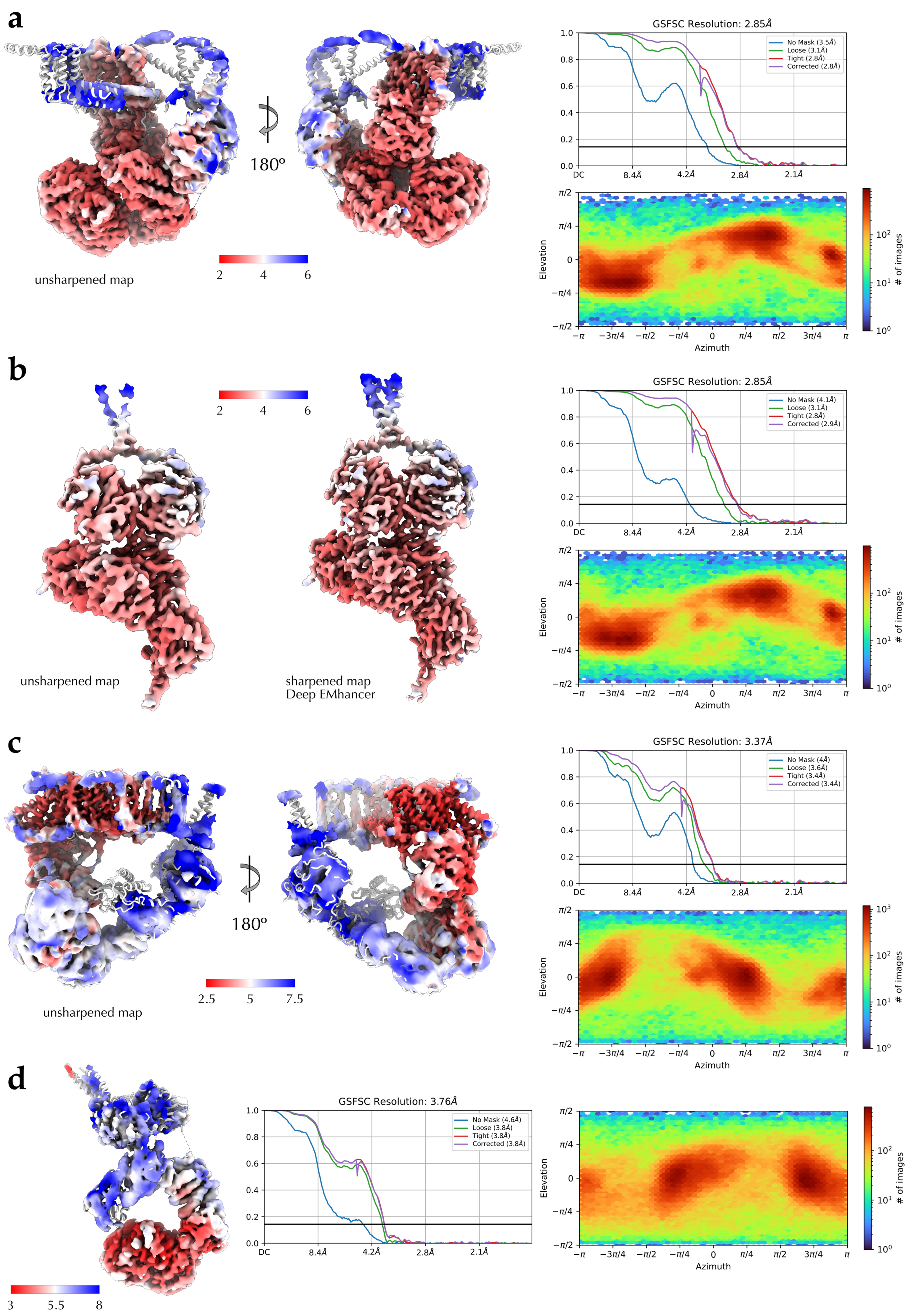


**Supplementary Fig. 5 Cryo-EM structures of the Bcs macrocomplex and the locally refined vestibule subcomplexes, supplemental information.** Local resolution surface mapping on the unsharpened or sharpened maps, GSFSC curves and viewing direction distributions are shown for each locally refined assembly.
